# B cell-stromal cell cross talk drives mesenteric lymph node eosinophilia during intestinal helminth infection

**DOI:** 10.1101/2023.10.28.564531

**Authors:** Emily Bessell, Rachel Finlay, Louisa K. James, Burkhard Ludewig, Nicola L. Harris, Matthew R. Hepworth, Lalit Kumar Dubey

## Abstract

Eosinophils are involved in host protection against multicellular organisms including helminths and often participate in regulating long-lasting humoral responses. However, their recruitment to the gut-draining mesenteric lymph node (mLN), where they support the development of the adaptive immune response is still elusive. Here, we demonstrate the mechanism underlying the recruitment of eosinophils to the murine mLN post gastrointestinal helminth infection. We found that mLN eosinophils accumulated at immune interactive sites such as the interfollicular and paracortical regions in an IL-4Rα-dependent manner and was directly associated with the reduced availability of stromal derived eosinophil chemoattractants. Using multiplex imaging we confirmed that eosinophils associate within a stromal niche containing Lyve1^+^ lymphatic vessels, ER-TR7^+^Pdpn^+^ FRCs, and extrafollicular CD138^+^ plasma cells. Experiments utilising complete and mixed bone marrow chimeras demonstrated that mice lacking IL-4Rα expression or LTβ expression selectively on B cells had diminished eosinophilia and reduced extrafollicular plasma cell numbers within the mLN. When co-cultured with LTβR activated FRCs, eosinophils gained an active phenotype with enhanced *Il1rl1* (ST2) receptor expression. LTβR ligation on FRCs resulted in enhanced IL-33 expression along with enrichment of distinct reactomes. Additionally, deletion of LTβR in FRCs reduced the homing capability of eosinophils to the mLN, confirming the significance of lymphotoxin signalling in granulocyte recruitment. Overall, these results highlight the previously unknown role of B cell-stromal cell crosstalk in driving mLN eosinophilia and their potential role in regulating the quality and magnitude of the humoral immune response generated within the mLN.

## Introduction

Lymph nodes (LNs) are highly organised structures essential for eliciting the adaptive immune response by facilitating immune cell interactions. The non-haematopoietic portion of LNs consist of fibroblastic and endothelial stromal cells which regulate and support immune cell migration and interaction within and to the LN (*1*). Various stromal subtypes including fibroblastic reticular cells (FRCs), lymphatic endothelial cells (LECs), and follicular dendritic cells (FDCs) have been implicated in regulating the recruitment, migration and localisation of B cells, T cells, and dendritic cells (DCs) within the LN to foster close interaction between the immune cells and facilitate the adaptive immune response (*2–5*). The recruitment of lymphocytes and DCs to the LN requires CCR7 expression and is mainly driven by CCL19, CCL21, and CXCL13 produced by FRC subsets (*3, 6–9*). In contrast, emerging evidence suggests that granulocytes home to the LN in a CCR7 independent manner to support lymphocyte function (*10, 11*). However, the association of lymphoid stromal cells with granulocytes such as eosinophils and their recruitment to distinct lymphoid niches during homeostasis and/or during inflammation is understudied.

Eosinophils are effector cells that mainly reside in the gastrointestinal tract and play a major role in homeostasis, as well as in settings of type-2 immunity and inflammation, including helminth infections and allergy (*12*). Eosinophils are terminally differentiated cells that originate in the bone marrow from granulocyte/macrophage progenitors (GMP) and are present at low levels in the blood and mucosal surfaces under homeostatic conditions, such as the intestinal lamina propria and lungs. They accumulate in larger numbers in inflamed or infected tissues via a type 2 cytokine mediated axis (*13*), and not only amplify the type 2 response by secreting cytokines IL-4, IL-5, and IL-13 (*14, 15*) but cause extensive tissue damage during worm expulsion due to degranulation; and as a result eosinophils are considered to be final stage effector cells. Alongside their effector functions, studies have shown that eosinophils have a major role in supporting the lymphoid adaptive immune response via B cell priming and survival (*16, 17*), antibody class-switching and maintaining gut homeostasis (*18*).

B cell follicle formation and B cell survival are commonly associated with lymphoid stromal cells, and we have previously shown that following infection with gastrointestinal helminth, *Heligmosomoides polygyrus (Hp), g*ut homeostasis is perturbed (*19*). The anti-helminth type-2 response within the draining mesenteric lymph node (mLN) is characterised by an accumulation of T-helper 2 (Th2) CD4^+^ T-cells, B cell follicle expansion, and extrafollicular B cell accumulation, which is supported by stromal subset expansion, proliferation, and remodelling (*20–23*). FRC crosstalk with B cells governs the overall lymphoid stromal expansion and chemokine secretion to support strategic immune cell positioning, yet the role of B cell-stromal cross-talk in driving the recruitment of eosinophils into the mLN is poorly understood (*9, 24*)

We and others have previously shown that following infection with *Hp,* IL-4, IL-5, and IL-13 were significantly increased in the mLN (*25*). IL-4, IL-5 and IL-13, secreted from Th2 cells and group 2 innate lymphoid cells (ILC2s) activate eosinophils and drive eosinophilia in infected or inflamed tissues by inducing the secretion of eosinophil-specific chemokines eotaxin-1 (CCL11), eotaxin-2 (CCL24), eotaxin-3 (CCL26) and RANTES (CCL5) from epithelial cells, macrophages, mononuclear cells and fibroblasts. However, the contribution of chemokine induced eosinophil recruitment into lymphoid niches is largely unexplored and little is known about which cellular sources provide these eosinophil chemoattractants.

Given the central role of stromal cells and B cells during the anti-helminth response, which overlaps with the role of eosinophils in type-2 immunity and maintenance of B cell function, we asked what role B cells and stromal cells might play in eosinophil recruitment to the mLN and how this might support humoral immunity to helminth infection (*24*). Here we have used a combination of bone marrow chimeras and advanced imaging methods, as well as flow cytometry and gene expression analysis to determine the mechanism by which eosinophils are recruited into the mLN post-gastrointestinal helminth infection. Using the model murine helminth, *Heligmosomoides polygyrus,* we found that eosinophils are recruited to the mLN in an interleukin-4 receptor α-chain (IL-4Rα) dependent manner. Our results highlight a novel role of IL-4Rα signalling during helminth infection which is important for B cell-stromal crosstalk leading to chemoattractant secretion by stromal cells that guides the recruitment, accumulation, and survival of eosinophils within the mLN.

## Results

### Mesenteric lymph node eosinophilia requires IL-4Rα during helminth infection

*Heligmosomoides polygyrus* (*Hp*) is a natural murine gastrointestinal helminth and serves as an excellent tool for studying type-2 immunity and as a model of chronic human helminth infection (*26*). Helminth infection induces eosinophilia into infected tissues, however, the presence of eosinophils in secondary lymphoid organs like the mLN is understudied. In order to evaluate mLN eosinophilia, we infected C57BL/6J mice with *Hp* and harvested the mLN from naïve and infected mice. Using flow cytometry we identified CD45^+^CD3^-^CD19^-^CD11c^-^ Ly6G^-^CD11b^+^Siglec-F^+^ cells as eosinophils (Fig. S1A) which were significantly increased within the mLN over the time course of infection (Fig. 1A-B, Fig. S1B). Both the percentage and absolute number of eosinophils within the mLN were significantly higher at 12- and 21 days post infection (dpi) compared to naïve mice (Fig. 1A-B). During *Hp* infection, a strong Th2 response is generated within the mLN, characterised by high levels of IL-4, IL-5, and IL-13 (*25*). As both IL-4 and IL-13 signal through IL-4Rα, which is key to anti-helminth immunity (*26*), we utilised IL-4Rα^-/-^ as well as mice in which IL-13 had been deleted by a reporter allele insertion (IL-13^gfp/gfp^ mice) to assess the role of these cytokine signalling pathways in mediating mLN eosinophilia. Following *Hp* infection, the entire chain of mLNs were harvested from naïve and infected WT, IL-4Rα^-/-^ and IL-13^gfp/gfp^ mice and analysed using flow cytometry. We observed a significant increase in the total weight and cellularity of the mLN post infection in WT but not in IL-4Rα^-/-^ mice (Fig. S1C-D), with the marked increase in eosinophil numbers seen in the infected WT mice ablated in infected IL-4Rα^-/-^ mice (Fig. 1C-F). However, depletion of IL-13 alone (IL-13^gfp/gfp^) was not sufficient to completely reduce the number of eosinophils following infection, suggesting IL-13 signalling is dispensable for the recruitment of eosinophils into the mLN (Fig. S1E-F). As eosinophilia into infected tissues is IL-5 dependent, we also analysed the number of eosinophils in the mLNs of infected mice where IL-5 had been depleted (IL-5^R5/R5^ mice) and found that these mice had a significant reduction of eosinophils post *Hp* infection (Fig. S1G-H), confirming IL-5 also plays a role in eosinophilia into the mLN, as expected. As the spatial organisation of the T cell and B cell zones following *Hp* infection is mediated by IL-4 signalling via IL-4Rα (*9*), we considered the possibility that the lack of eosinophilia in IL-4Rα^-/-^ mice may be a result of differential stromal remodelling (*21*). To investigate this in detail, we conducted high-resolution multiplex imaging of WT and IL-4Rα^-/-^ mLN post *Hp* infection. These imaging studies revealed that WT infected mice had an increased accumulation of Siglec-F^+^ eosinophils in the cortical and FRC rich paracortical regions (Fig. 1G). Interestingly, in IL-4Rα^-/-^ mice there were few Siglec-F^+^ cells in the paracortical T cell zone (Fig. 1H) with some accumulation of eosinophils in the medullary region. Collectively, these observations indicated that Siglec-F^+^ eosinophils are recruited to and localise in distinct regions of mLN following helminth infection in an IL-4Rα dependent manner.

**Figure 1:**
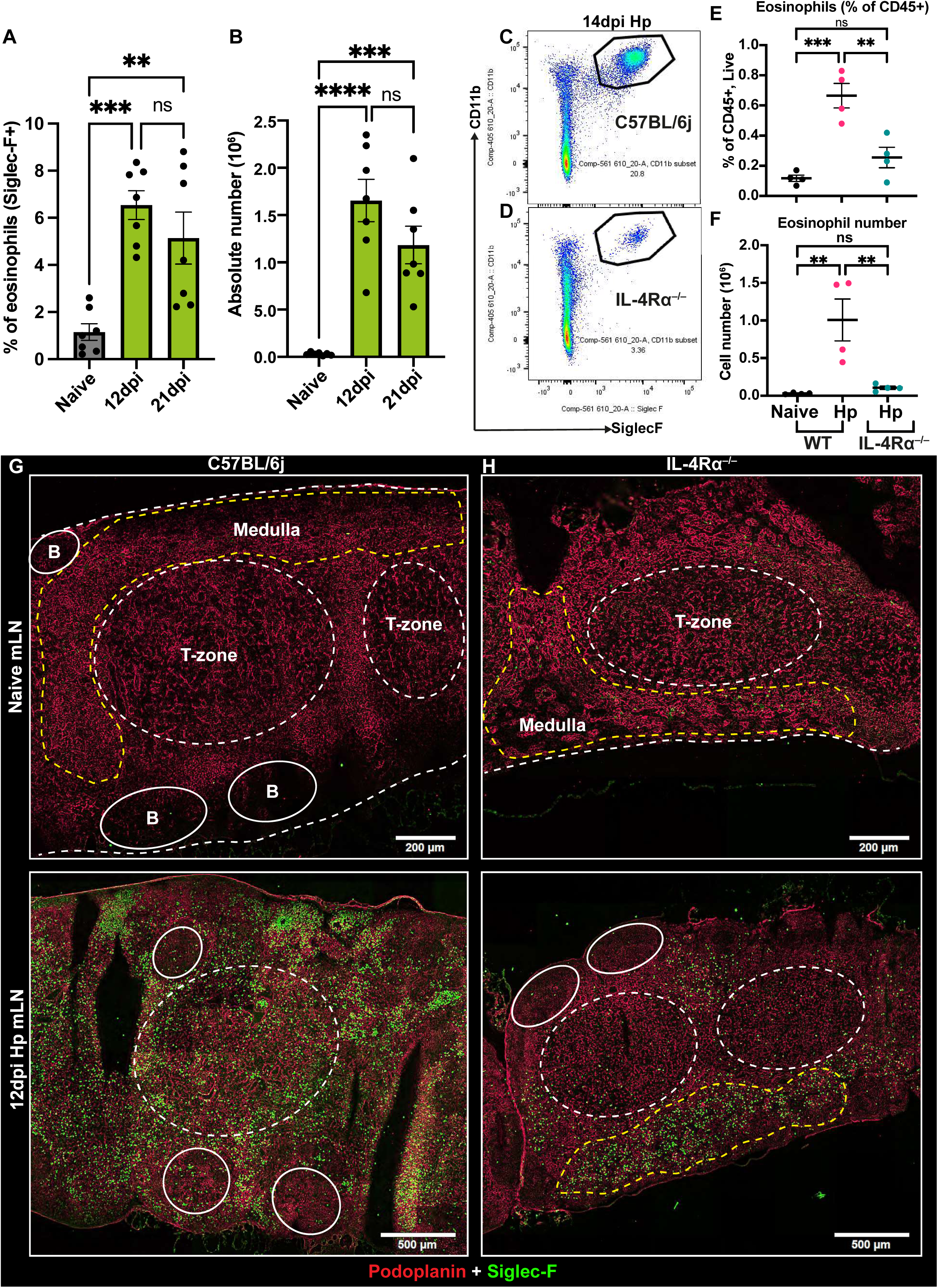
Mesenteric lymph node eosinophilia requires IL-4Rα. C57BL/6J (WT) mice were infected with *Hp* and the entire chain of the mLN were collected at 0 (naïve), 12, and 21 dpi and analysed by flow cytometry. **(A)** Percentage and **(B)** total number of CD11b^+^Siglec-F^+^ eosinophils within the mLN. Data is pooled from 2 different independent experiments. Data represents mean ± SEM with n ≥3-4 mice per group. \**P* < 0.05, \*\**P* < 0.01, \*\*\**P* < 0.001, and \*\*\*\**P* < 0.0001 (ANOVA, Bonferroni’s multiple comparison test). Pseudocolour dot plots showing eosinophils within the mLN of **(C)** WT and **(D)** IL-4Ra knockout (IL-4Rα**^−/−^**) mice post *Hp* infection. **(E)** Percentage of eosinophils gated on CD45^+^ live cells and **(F)** absolute number of Siglec-F^+^ eosinophils within the mLN of WT and IL-4Rα**^−/−^** mice. Whole mLN cryosections for **(G)** WT and **(H)** IL-4Rα**^−/−^** showing combined staining for the eosinophils (Siglec-F^+^, green) and stromal cells (Pdpn^+^, red). Scale bars = 200 μm and 500 μm. Images are from individual mice that are representative of ≥4 different experiments with n ≥2-3 mice/group/time point. The T zone (white dotted circle), B zone (solid circle), and medulla (yellow dotted line) is highlighted to identify the sub-regional organisation of mLN.

### Interfollicular accumulation of eosinophils requires IL-4Rα

Next, we used multiplex imaging to extend these observations by taking advantage of the fact that *Hp* infection induces stromal remodelling and recruits immune cells to immune interactive sites like the T/B border and interfollicular region (*9*). mLNs were harvested from WT and IL-4Rα^-/-^ mice at 0-(naïve) and 21 dpi and stained for LECs (Lyve1^+^) and eosinophils (Siglec-F^+^). LECs are key players in lymphocyte migration and during *Hp* infection expand throughout the LN to support B cell follicle formation and T cell-DC interaction (*27*). Lyve1 staining was performed to define and identify cortical, medullary, and paracortical segregation. The naïve mLN from both WT and IL-4Rα^-/-^ mice showed no sign of eosinophilia (Fig. S2A-B). In WT mice, eosinophils were recruited to the paracortical region (Fig. 2C, inset 1) and the IFR (Fig. 2D, inset 2) at 21 dpi, whereas in IL-4Rα^-/-^ mice, fewer eosinophils were recruited into the IFR and/or the T/B border (Fig. 2E). This suggested that IL-4Rα supported the migration of eosinophils into mLN immune interactive sites. High-resolution images of 21 dpi mLN from WT and IL-4Rα^-/-^ mice showed a close association of lymphatic vessels and follicular B cell clusters alongside eosinophils (Fig. S2C) suggesting that eosinophils access the cortical lymphatic sinuses, which are known to serve as egress sites for immune cells in close association with the B cell follicle (*28, 29*) and potentially support B cell function by providing survival factors and support antibody isotype class switching (*16, 30*). To further confirm the eosinophil association with stromal cells, 21 dpi mLN from WT and IL-4Rα^-/-^ mice were subjected to deep tissue imaging. Staining of vibratome sections further confirmed the enhanced eosinophilia in WT mice compared to IL-4Rα^-/-^ mice confirming the 2D histological findings (Movies S1-2). The observed close association of eosinophils with both FRCs and LECs suggests a potential role of stromal cells in mLN eosinophilia. Collectively, these observations highlight the previously unknown association of eosinophils with lymphoid stromal cells and suggests that IL-4Rα is key to their accumulation in immunological niches.

**Figure 2:**
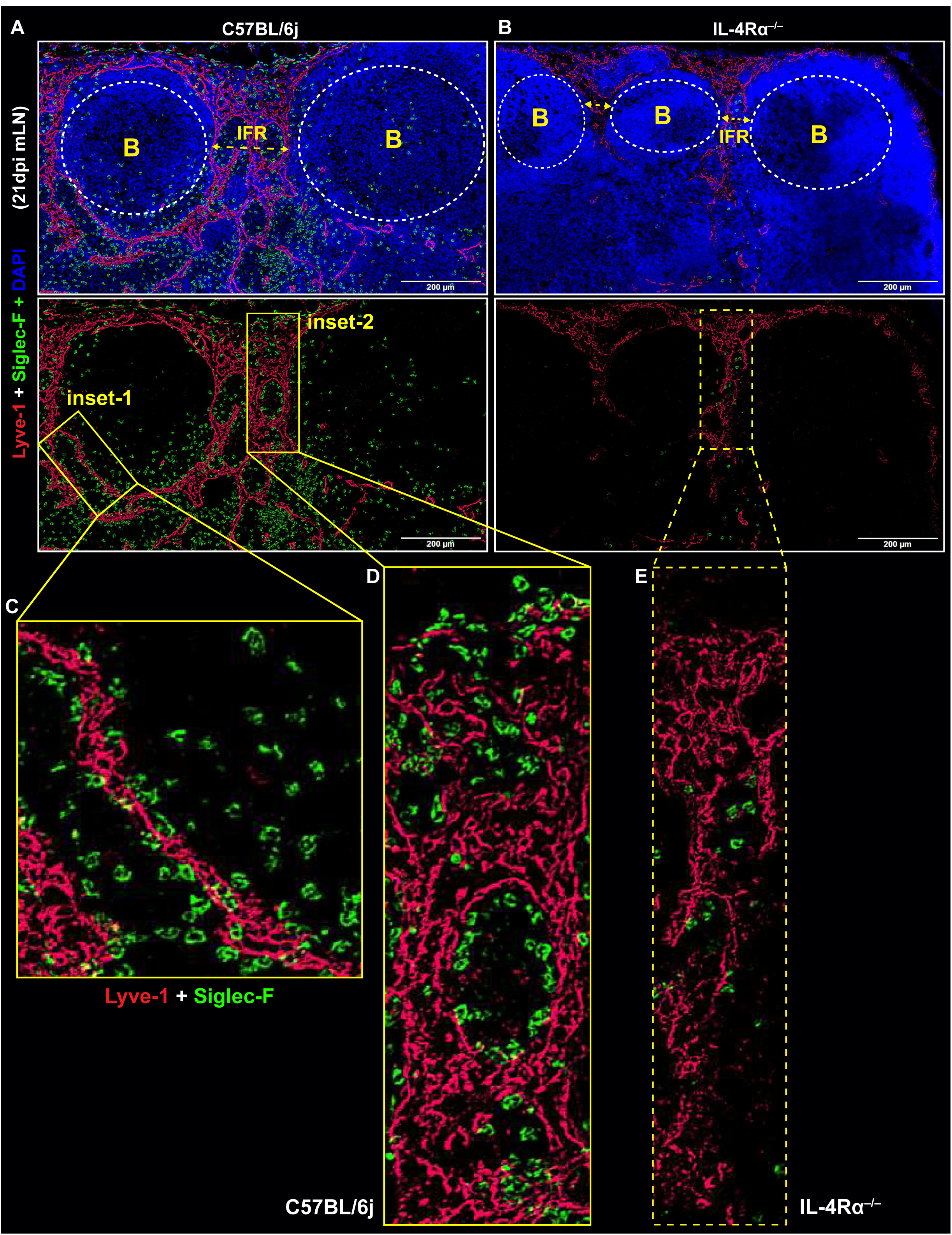
Interfollicular accumulation of eosinophils requires IL-4Rα. WT and IL-4Ra**^−/−^**mice were infected with *Hp* and the mLN was collected at 21 dpi. mLN cryosections showing combined staining for DAPI (blue), eosinophils (Siglec-F^+^, green), and lymphatic endothelial cells (Lyve1^+^, red) cells in **(A)** WT and **(B)** IL-4Rα**^−/−^** mice post *Hp* infection. The yellow inset-1 and inset-2 highlight the eosinophil accumulation at the paracortical region, T/B border, and interfollicular regions (IFR) respectively. Eosinophil accumulation at **(C)** paracortical and T/B border **(D)** IFR in WT mice and **(E)** IFR in IL-4Rα**^−/−^** mice. Scale bar = 200 μm. Images are representative of ≥5 different experiments with n ≥2-3 mice/group/time point.

### Non-haematopoietic stromal cells provide eosinophil chemoattractant

Given the increase of eosinophils in the mLN post infection, translocation to distinct lymphoid niches, and association with stromal cells (Figs. 1 and 2), we speculated that chemokine signals may arise from the mLN stroma to support eosinophil recruitment. To evaluate this hypothesis, the total CD45^-^ stromal cells from naïve and infected WT and IL-4Rα^-/-^ mice were isolated for gene expression analysis. The three key murine eosinophil chemoattractants, RANTES (CCL5), eotaxin-1 (CCL11), and eotaxin-2 (CCL24) were evaluated for their relative expression. All three genes were expressed by the CD45^-^ stromal fractions (Fig. 3A-C). A strong upregulation of *Ccl24* was observed during the early phase of infection in WT mice in an IL-4Rα dependent manner, which remained elevated at later time points (Fig. 3C), whereas an increase in *Ccl11* was observed during the later phase of infection, i.e., 12 dpi (Fig. 3B). Interestingly, analysis of the key stromal subsets using RNA-seq database Immgen.org, confirmed the differential expression of eosinophil chemoattractants by the key stromal subsets found in secondary lymphoid organs (Fig. 3D). All subsets express some CCL5 while FRCs had the highest expression of *Ccl11* and LECs expressed the most *Ccl24*. Considering an enhanced *Ccl24* expression in the stromal fraction post infection, we further validated these findings using immunofluorescence microscopy with naïve and infected mice mLN cryosections. mLN sections were stained with Lyve1, podoplanin (Pdpn) and CCL24. Interestingly, there was little to no detectable expression of CCL24 in naïve mLN (Fig. 3E). However, post *Hp* infection at 12 and 21 dpi, a significant increase in CCL24 expression was observed. Closer analysis revealed that Lyve1^+^ LECs and not Pdpn^+^ FRCs were the main source of CCL24 post-infection (Fig. 3F-G). These data suggested that the lymphatic endothelium is the principal cellular source of CCL24 during infection and provides an explanation for the preferential localisation of eosinophils in proximity to the lymphatic vessels. Overall, these findings highlight the role of IL-4Rα driven chemoattractant expression by stromal subsets within the mLN which in turn regulate the eosinophil recruitment and localisation.

**Figure 3:**
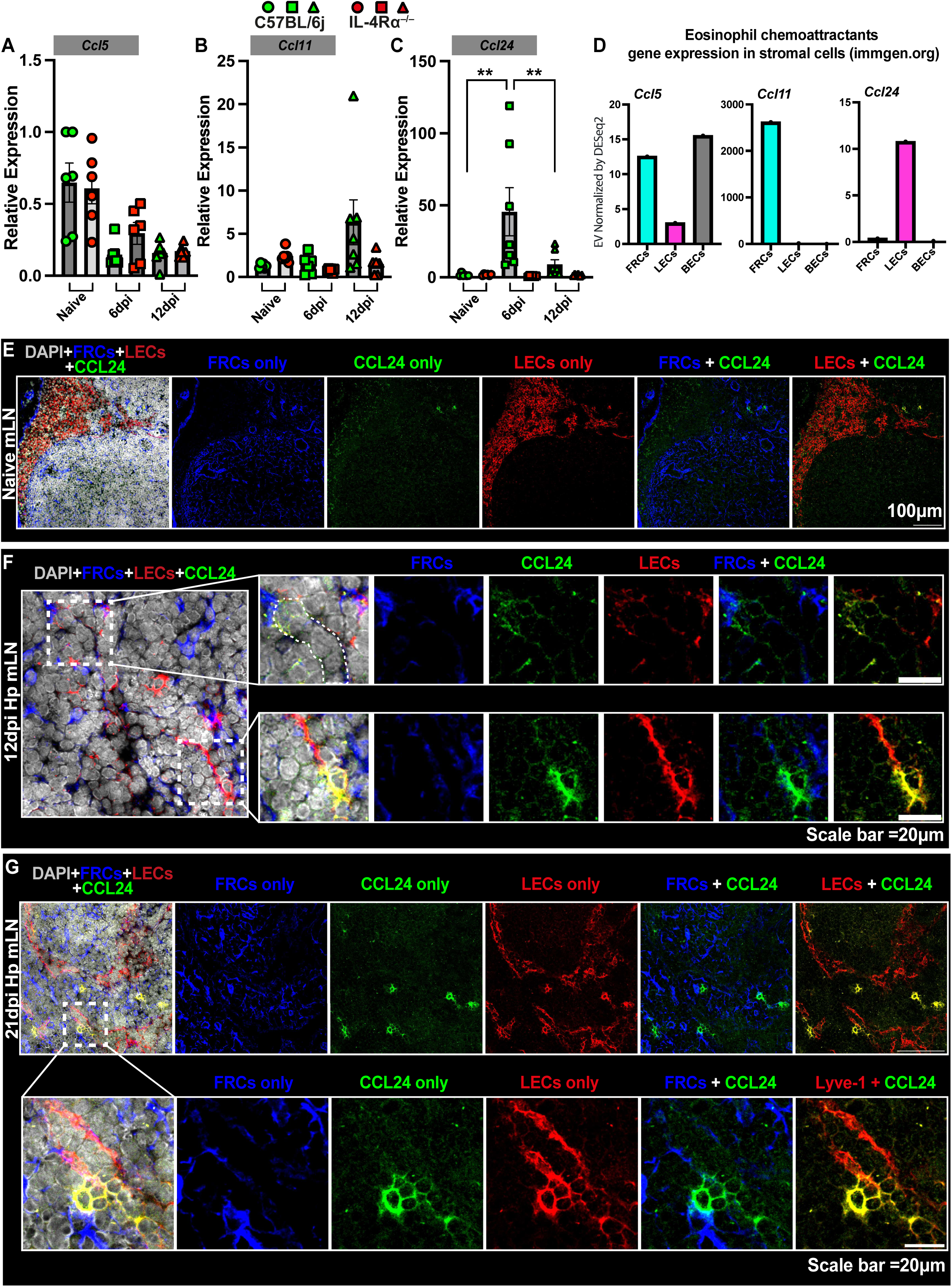
Non-haematopoietic stromal cells provide eosinophil chemoattractant to support mLN eosinophilia. C57BL/6J (WT, green histogram) and IL-4Rα**^−/−^** (red histogram) mice were infected with *Hp,* and the entire chain of the mLN was collected at 0 (naïve), 6 dpi, and 12 dpi. The total CD45^-^ stromal fraction were isolated and used for gene expression analysis. The relative expression of the major eosinophil chemoattractants, **(A)** *Ccl5*, **(B)** *Ccl11,* and **(C)** *Ccl24* were analysed using real-time PCR. Pooled data from two independent experiments with n ≥3 mice per group are shown as mean ± SEM. *P < 0.05, **P < 0.01, ***P < 0.001, and ****P < 0.0001 (ANOVA, Bonferroni’s multiple comparison test). **(D)** Analysis of public gene expression database (Immgen.org) for eosinophil chemoattractant genes within the three most abundant stromal populations. mLN cryosections showing combined staining for DAPI (grey), ER-TR7^+^ FRCs (blue), CCL24^+^ (green), and Lyve1^+^ lymphatic endothelial cells (LECs, red) in WT mice during **(E)** naïve, **(F)** 12 dpi and **(G)** 21 dpi *Hp* infection. **(F-G)** White dotted insets show the enhanced CCL24 expression on Lyve1+ vessels. Scale bar= 100 μm and 20 μm.

### IL-4R**α** dependent cortical/medullary association of eosinophils with extrafollicular plasma cells

Following the observation that eosinophils localise within close vicinity of B cell follicles and lymphatics, we hypothesised that they would also be closely associated with mLN extrafollicular B cells and plasma cells. mLN sections from naïve and infected WT and IL-4Rα^-/-^ mice were analysed immunohistochemically to confirm the localisation of CD138^+^ plasma cells with the lymphatic sinuses. Immunohistochemical staining of mLN serial sections showed that following infection there is an enhanced extrafollicular B cell response with a significant expansion of Lyve1^+^ sinuses, whereas IL-4Rα^-/-^ infected mice failed to mount such response (Fig. S3A-B). We observed that in the cortical/medullary and paracortical regions of the mLN in WT naïve mice there was little to no presence of eosinophils (Siglec-F^+^) and plasma cells (CD138^+^) (Fig. 4A, insets i, ii). Additionally, in the paracortical region, we observed no lymphatics (Lyve1^+^), which correlated with the lack of eosinophils and plasma cells in this region. Comparatively, in the WT 21 dpi mice we observed a significant increase in both eosinophils and plasma cells in the cortical/medullary and paracortical regions (Figure 4B, insets iii, iv) a feature not evident in mice lacking IL-4Rα (Fig. 4C, insets v, vi). Quantification of immunofluorescence images, highlighting the area occupied by plasma cells and eosinophils, complemented the immunohistochemical staining showing that eosinophils and plasma cells co-existed within the mLN (Fig. 4D-E). Additionally, analysis by flow cytometry highlighted that expansion of the plasma cell population following *Hp* infection is IL-4Rα dependent, with a significant reduction in plasma cell numbers in IL-4Rα^-/-^ mice at 12 and 21 dpi compared to WT mice (Fig. 4F-G). These data suggest that following helminth infection, IL-4Rα driven expansion of the lymphatic network supported eosinophil and plasma cell co-localisation in the cortical/medullary and paracortical regions, which has the potential support the extrafollicular B cell response, which is key to anti-helminth antibody production (*20, 21, 31*).

**Figure 4:**
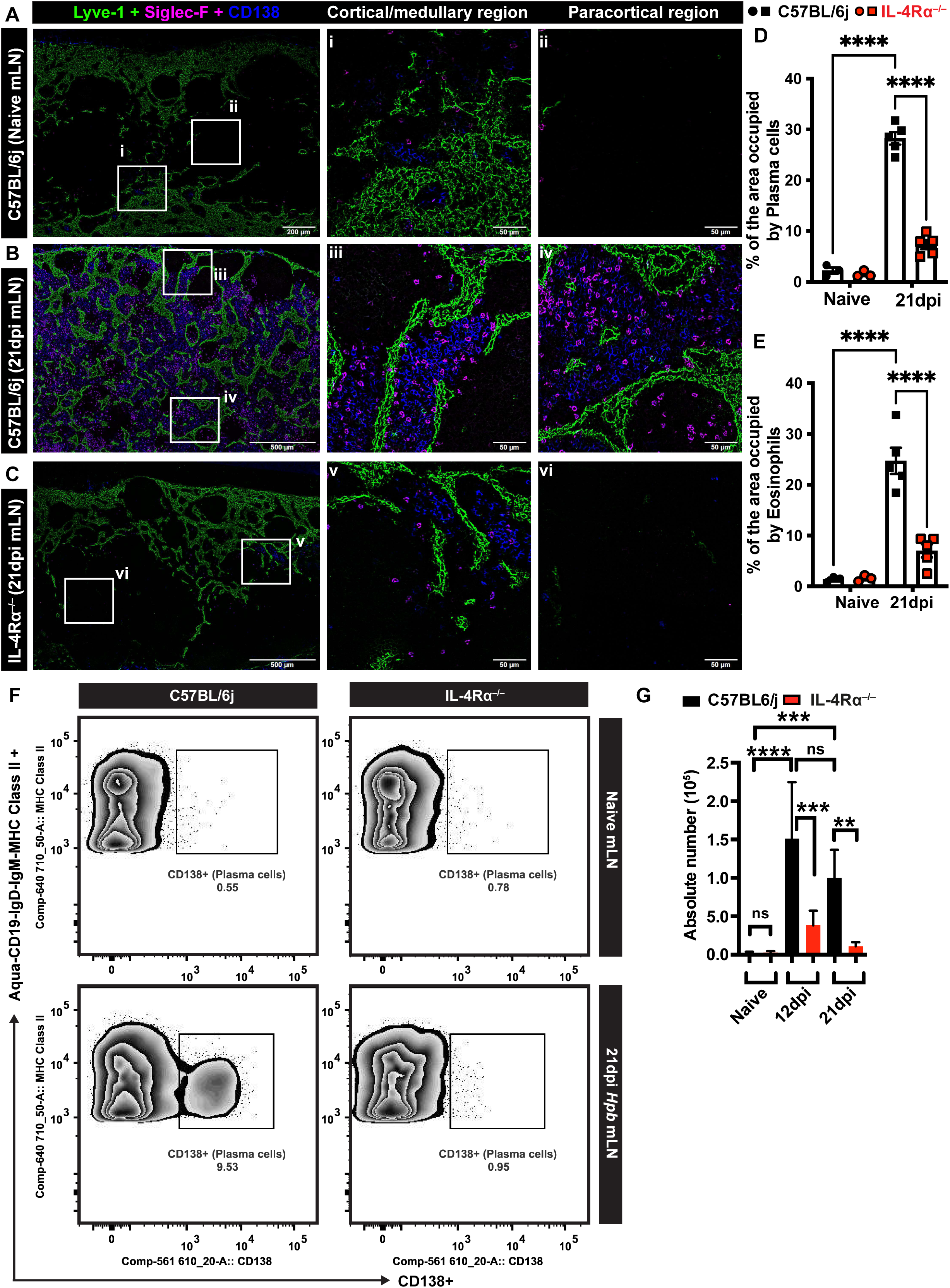
IL-4Rα dependent cortical-medullary association of eosinophils with extrafollicular plasma cells. C57BL/6J (black histogram) and IL-4Rα **^−/−^** (red histogram) mice were infected with *Hp* and the entire chain of the mLN was collected at 0 (naïve) and 21 dpi. **(A-B)** WT and **(C)** IL-4Rα**^−/−^** mLN cryosections showing combined staining for the CD138^+^ plasma cells (blue), Siglec-F^+^ eosinophils (magenta), and Lyve1^+^ LECs (green). The white insets i, iii, and v highlight the cortical/medullary region, and insets ii, iv, and vi highlight the paracortical region. Scale bar = 200 μm (naïve mLN) and 500 μm (Hp mLN). Scale bar in the insets is 50 μm. Images are representative of ≥3 different experiments with n ≥2-3 mice/group/time point. Quantification of WT and IL-4Rα**^−/−^** mice immunofluorescence images from the mLN showing % area coverage by **(D)** CD138^+^ cells and **(E)** Siglec-F^+^ cells. **(F)** Dot plots show the plasma cell population in the WT and IL-4Rα^-/-^ naïve and infected mLN. **(G)** Absolute number of CD138^+^ plasma cells in the mLN of naïve, 12 dpi, and 21 dpi WT and IL-4Rα^-/-^ mice. Data represents mean ± SEM and are representative of two independent experiments with n ≥2-3 mice per infected group. \**P* < 0.05, \*\**P* < 0.01, \*\*\**P* < 0.001 and \*\*\*\**P* < 0.0001 (non-parametric Mann Whitney T test).

### IL-4R**α** expressing B cells drive mLN paracortical eosinophilia

We next investigated whether IL-4Rα expression on non-haematopoietic stromal cells (CD45^-^) or haematopoietic cells (CD45^+^) were important for mLN eosinophilia. To assess this, we performed complete bone marrow chimeras where IL-4Rα^-/-^ recipient mice were given bone marrow from WT donor mice to generate mice lacking IL-4Rα on stromal cells (Fig. 5A). In parallel, a cohort of WT mice (recipient) with IL-4Rα^-/-^ mouse bone marrow (donor) were generated to analyse mice lacking IL-4Rα on CD45^+^ haematopoietic cells (Fig. 5B). Post chimerism, mice were infected with *Hp* and the whole mLN chain was harvested from mice at 21 dpi as well as from naïve mice. In both bone marrow chimera models, there was no eosinophilia (Siglec-F^+^) in the paracortical region in naïve mice (Fig. S4A-D). At 21 dpi, eosinophilia was observed in the mice lacking IL-4Rα on stromal cells (Fig. 5C), but no eosinophilia in mice lacking IL-4Rα on haematopoietic (CD45^+^) cells (Fig. 5D), indicating IL-4Rα expression on haematopoietic cells is required for the recruitment of eosinophils to the paracortical region of the mLN.

**Figure 5:**
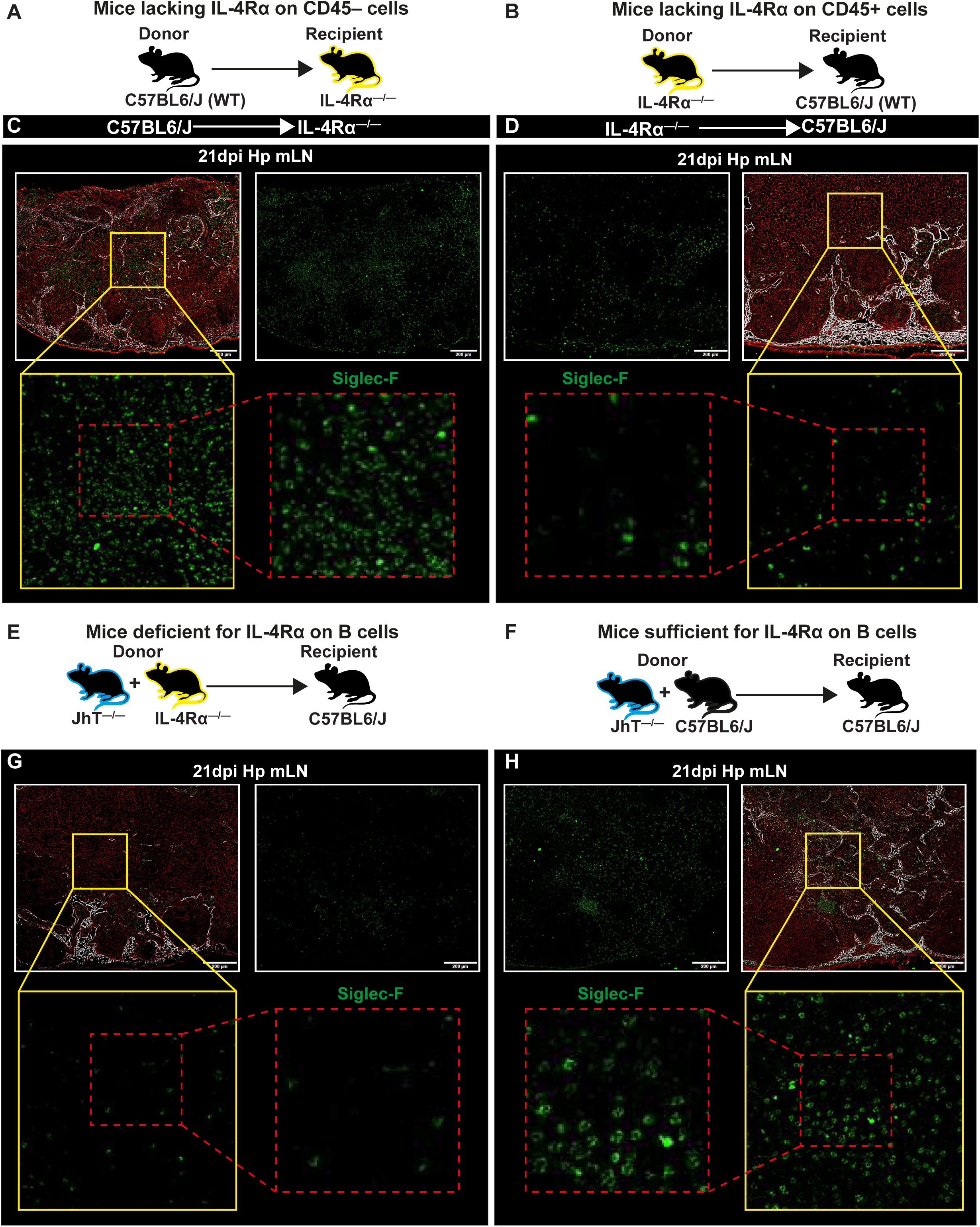
IL-4Rα expressing B cells drive mLN paracortical eosinophilia. Complete bone marrow chimera mice (donor strain/recipient strain) showing mLN eosinophilia during helminth infection. **(A)** IL-4Rα^−/−^ mice received C57BL/6J (WT) bone marrow and **(B)** C57BL/6J mice received IL-4Rα^−/−^ mice bone marrow. The resulting animals lacked IL-4Rα on **(A)** CD45^-^ and **(B)** CD45^+^ cells. **(C-D)** mLN cryosections from chimeric infected mice showing immunofluorescence image staining for Lyve1^+^ (white), Pdpn^+^ (red), and Siglec-F^+^ (green) are shown. Mixed bone marrow chimeras were generated using lethally irradiated wildtype (WT) recipients reconstituted with € bone marrow cells from B cell deficient (Jht^−/−^) mice mixed with bone marrow cells from IL-4Rα^−/−^ mice (Jht^−/−^ + IL-4Rα^−/−^). **(F)** Control mice received B cell deficient bone marrow mixed with WT cells (Jht^−/−^ + WT). All mice were infected with *Hp* and the mLN was collected at day 0 (naïve) and 21 dpi for analysis. **(G-H)** mLN cryosections from Jht^−/−^ + IL-4Rα^−/−^ and control mixed bone marrow chimeric infected mice showing immunofluorescence image staining for Lyve1^+^ (white), Pdpn^+^ (red) and Siglec-F^+^ (green). The magnified view of the paracortical region (yellow and red insets) from 21 dpi mLN highlighting eosinophilia is shown for both groups in a side-by-side comparison. Scale bar = 200 µm. The red insets show a higher magnification of Siglec-F^+^ cells (green) within the paracortical region. The images are from representative mice and from two independent experiments with n ≥2-4 mice per group.

Having confirmed that IL-4Rα expression on haematopoietic cells is important for the recruitment of eosinophils we next investigated whether IL-4Rα expression on B cells was important for the recruitment of eosinophils into the mLN. To determine this, a mixed bone marrow chimerism approach was used where WT mice were lethally irradiated and reconstituted with bone marrow from B cell deficient mice (Jht^-/-^) and IL-4Rα^-/-^ mice to produce resultant mice that lacked IL-4Rα expression on B cells (Jht^-/-^ + IL-4Rα^-/-^, Fig. 5E). As a control, WT mice received bone marrow from B cell deficient mice (Jht^-/-^) and WT mice (Jht^-/-^ + WT, Fig. 5F). We observed no apparent difference in the numbers of eosinophils in mice with IL-4Rα deficient B cells and control mice under steady state (Fig. S4E). At 21 dpi with *Hp*, mice lacking IL-4Rα expression on B cells failed to remodel the IFR as well as the paracortical region (*21*) which corresponded with reduced eosinophil numbers compared to mice that are sufficient for IL-4Rα on B cells (Fig. 5G, H). These results confirmed that IL-4Rα expressing B cells are important for the recruitment of eosinophils into the paracortical region of the mLN.

### Lymphotoxin expressing B cells govern mLN eosinophilia

Lymphotoxin-beta (LTβ) is a key to lymphoid organ development and architecture and regulates interactions between stromal cells and lymphocytes (*32–34*). We have previously shown that IL-4Rα signalling on B cells drives LTα_1_β_2_ (lymphotoxin) upregulation, which then interacts with LTβR expressing stromal cells to govern lymphoid remodelling during a helminth infection (*9*). Having confirmed the role of IL-4Rα on B cells for driving eosinophilia into the mLN, we hypothesised that IL-4Rα driven lymphotoxin expression on B cells might be required for eosinophil recruitment to the mLN. To address the role of lymphotoxin expressing B cells in driving mLN eosinophilia, we used a mixed bone marrow chimera approach. WT mice were lethally irradiated and reconstituted with bone marrow from B cell deficient (Jht^-/-^, Fig. 6A) or T cell deficient (TCRβδ^−/−^, Fig. 6C) mice and mixed with bone marrow from LTβ^-/-^ mice. The resultant mice lacked the expression of LTβ on either B cells (Jht^-/-^ + LTβ^-/-^, Fig. 6A) or on T cells (TCRβδ^−/−^ + LTβ^-/-^, Fig. 6C). Control mixed bone marrow chimeras were also generated using WT donors alongside LTβ^-/-^ (Jht^-/-^ + WT, Fig. 6B) and TCRβδ^−/−^ (TCRβδ^−/−^ + WT, Fig. 6D). The recipient mice were infected with *Hp* and the mLN was collected at 21 dpi. Under steady state conditions, we observed no apparent difference in terms of lymphoid organisation and numbers of eosinophils in LTβ deficient B cells and LTβ deficient T cells as well as in control mice (Fig. S5A-D). Furthermore, no paracortical eosinophilia was observed in these animals. However, post *Hp* infection, mice that were selectively deficient for LTβ expression on B cells failed to expand the lymphatics network and were unable to recruit eosinophils into the interfollicular and paracortical regions of the mLN (Fig. 6A). Higher magnification images further confirmed the reduced eosinophil association with cortical lymphatics (Fig. 6E, white dotted insets). Comparatively, mice sufficient for LTβ on B cells (Fig. 6B, F) were able to recruit eosinophils to the mLN and have an expanded lymphatics network following infection. Mice that were LTβ deficient or sufficient on T cells (Fig. 6C-D, G-H) had comparable eosinophilia in the mLN following infection. Eosinophils were seen in close association with Lyve1^+^ lymphatic endothelial cells (Fig. 6A-B) confirming that B cell crosstalk with stromal cells yields a favourable niche that supports eosinophil recruitment and interaction with stromal and immune cells within the mLN. Furthermore, these results confirmed that LTβ expression on B cells and not T cells in response to *Hp* infection is crucial for the recruitment of eosinophils into the follicle-proximal regions of the mLN by supporting the expansion of the stromal network.

**Figure 6:**
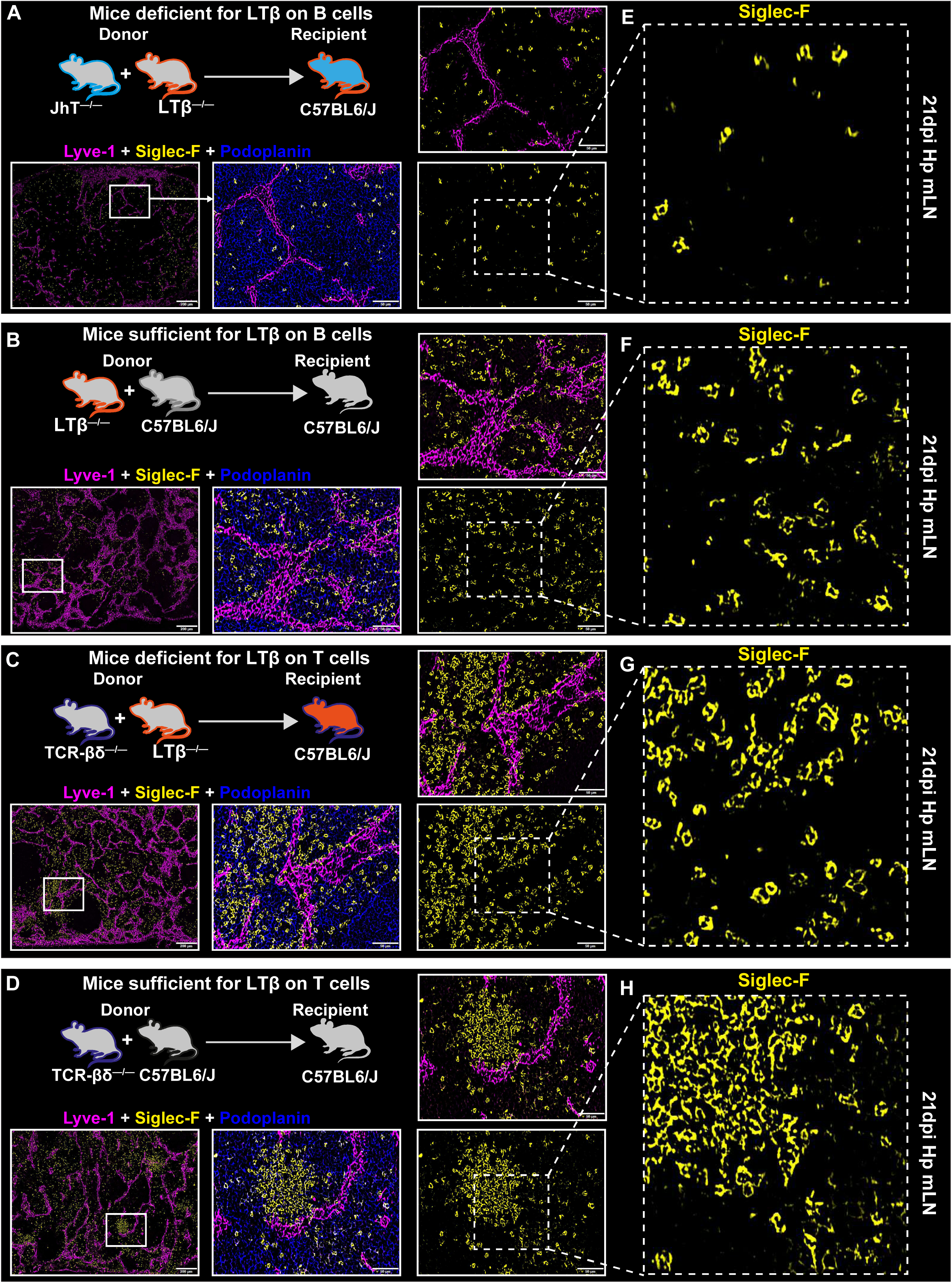
Lymphotoxin expressing B cells govern mLN eosinophilia. Mixed bone marrow chimeras were generated as described in the methods section. Chimeric mice lacking lymphotoxin expression distinctly on either **(A)** B cells (Jht^−/−^ + LTβ^−/−^) or **(C)** T cells (TCRβδ^−/−^ + LTβ^−/−^) were compared to respective control mice having **(B)** B-cell or **(D)** T-cell sufficient for lymphotoxin, Jht^−/−^ + WT and TCRβδ^−/−^ + WT, respectively. All chimeric mice were infected with *Hp* and the mLN was collected at day 21 dpi. Infected chimeric mice lacking lymphotoxin on **(E)** B cells (Jht^−/−^ + LTβ^−/−^) and their respective **(F)** controls (Jht^−/−^ + WT) showing immunofluorescence staining for Pdpn^+^ (blue), Siglec-F^+^ (yellow) and Lyve1^+^ (magenta). Infected chimeric mice lacking **(G)** lymphotoxin on T cells (TCRβδ^−/−^ + LTβ^−/−^) and respective **(H)** controls (TCRβδ^−/−^ + WT) showing eosinophilia (yellow cells) along the LECs network (magenta). The white inset shows a higher magnified view of Siglec-F^+^ cells (yellow) in close vicinity to the interfollicular region. Scale bar = 200 μm and 50 µm.

### LT**β**R activated FRCs enhances eosinophil activation and migratory capacity

Having established that eosinophils are present at the IFR and paracortical region were also in close proximity with B cells on the stromal scaffold, we hypothesised that eosinophil activation and gene expression were linked to their interaction with the activated stroma. To directly assess this hypothesis, we ligated LTβR on cultured mLN FRCs using a LTβR agonist antibody (Clone 4H8WH2) (Fig. 7A, condition 1); also considering eosinophils have long been associated with providing B cell proliferative and survival factors (*30*) we mimicked a cell-based activation model using purified naïve B cells (Fig.7A, condition 2). Both treatments resulted in an activated FRC phenotype and were later used for co-culturing with bone marrow-derived eosinophils. After 8 hours of co-culture, the eosinophils were purified, and bulk RNA sequencing was performed to identify global changes in the transcriptomes of eosinophils post co-culture with the activated FRCs. We identified that eosinophils co-cultured with B cell activated FRCs had a greater enrichment of *MhcII*, *Ccr7*, *Il6* and *Il1*β, (Fig. 7B-E), suggesting the interaction of B cells with FRCs provides alternative factors that enhance eosinophil activation as well as B cell priming capabilities. Furthermore, extracellular activation of eosinophils is regulated through the NF-κB signalling pathways (*35*), so we performed subsequent analysis to assess the expression of NF-κB signalling components: *Nfkbie, Relb, Nfkb1, Nfkbib, Rela, Nfkb2* and *Nfkbiz* in eosinophils obtained from the co-culture. In both conditions, comparable NF-κB activation was observed (Fig. 7F-G). We further analysed various known functional modules, including adhesion, tissue repair and degranulation (Fig. 7H-J), as well as cytokines, chemokines, and Ig-receptors (Fig. S6A-C), which are important for eosinophil function, and the gene expressed remained mostly conserved between the two groups. Analysis of cytokine/chemokine receptors in eosinophils showed an enrichment of the ST2 (*Il1rl1*) receptor along with *Il5ra* and *Tnfrsf1a* (target gene for STAT 3) (Fig. 7K). We next performed STRING analysis to find putative direct and indirect interactions on these genes. The STRING analysis yielded a total of 31 biological processes based on gene ontology (GO) (Table S1). We focused on proteins relating to the regulation of IL-5 production (GO:0032674) considering IL-5 is one of the key survival factors for eosinophils. Out of 22 known proteins within the network (Fig. S6D), STRING analysis predicted co-expression and/or co-occurrence between the ST2 receptor (*Il1rl1*) and *Il5ra* within our network which could be linked to receiving survival signals from both IL-5 and IL-33 (Fig. 7L). The wider network representation further highlights a distinct cluster of proteins where eosinophil peroxidase (Epx) was also associated with IL5ra and IL-33 (Fig. S6D), suggesting IL-33 may also be involved with eosinophil immune functions. Overall, these results highlight a previously unidentified role of activated FRCs and the niches they create by interacting with B cells within the mLN, which in turn can modulate the eosinophil activation and gene expression.

**Figure 7:**
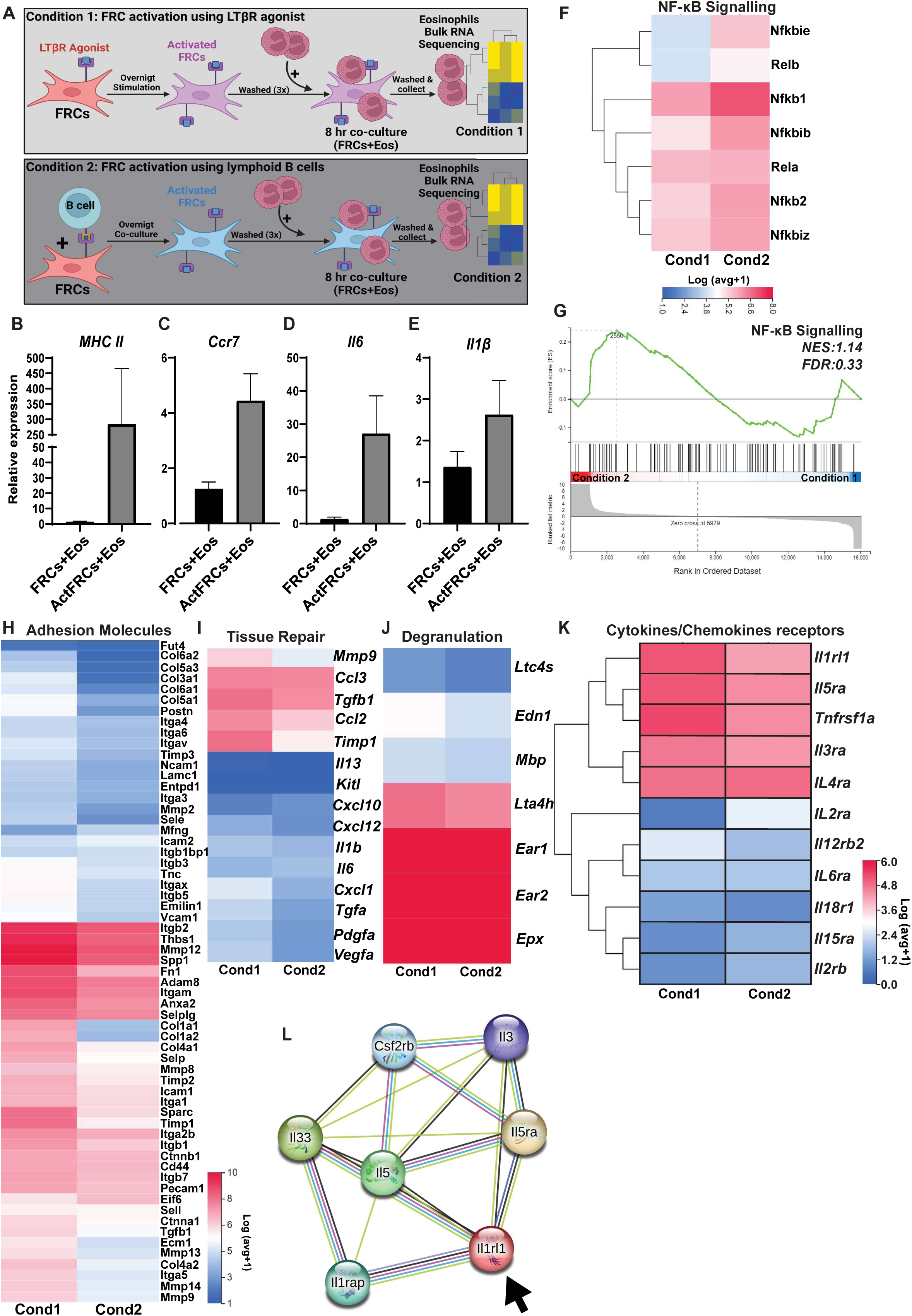
LTβR activated FRCs enhances eosinophil activation and survival. LN derived stromal cells and bone marrow-derived eosinophils were generated *in vitro* as described in the methods section. FRCs were activated with **(A)** LTβR agonist antibody (2 µg/mL) (condition 1) or with naïve B cells (condition 2). Eosinophils co-cultured with B cell-activated stromal cells were analysed for **(B)** *MhcII*, **(C)** *Ccr7*, **(D)** *Il6*, and **(E)** *Il1*β gene expression. **(F)** Heatmap and **(G)** Gene set enrichment analysis (GSEA) plot showing expression of genes linked to NFκB signalling. Heatmaps showing genes relating to **(H)** tissue repair, **(I)** degranulation, **(J)** adhesion molecules and **(K)** cytokine receptors. **(L)** STRING analysis was performed on cytokine receptors, STRING network showing cytokine receptors within the regulation of IL-5 production gene ontology pathway (GO:0032674). Known interactions are indicated by cyan (curated database) and magenta (experimentally determined), and predicted interactions are indicated by black (co-expression), green (textmining) dark blue (gene co-occurrence), and lilac (protein homology). The data is representative of at least 1 independent experiment and is presented as mean ± SEM. One-way ANOVA tests were used to denote the significance - *p<0.05, **p<0.01 ***p<0.001, ****p<0.0001, ns p>0.05.

### Loss of lymphotoxin-beta receptor (LT**β**R) on CCL19^cre+^ FRCs attenuates eosinophilia

We have previously established that lymphotoxin expressing B cells interact with LTβR expressing CCL19^+^ stromal cells to govern the expansion and remodelling of mLN niches and eosinophilia (Fig.4-6). We further validated these findings using our co-culture system where FRCs were activated through LTβR signalling. Ligation of LTβR using an agonist antibody enhanced the IL-33 expression in FRCs which was further enhanced by lymphotoxin-expressing B cells (Fig.8A). The activated phenotype of FRCs was confirmed by analysing ICAM-1 expression (Fig.8B) which is in line with previous reports (*36*). Since we have shown that LTβR activated stromal cells are required for mLN eosinophilia and FRC-derived IL-33 may play a role in eosinophil function via the ST2 receptor, we further evaluated how IL-33 influenced eosinophil gene expression. To explore how stromal-derived IL-33 influenced the gene expression profile of eosinophils, we performed bulk RNA sequencing analysis by utilising datasets of known IL-33 induced genes in eosinophils (GSE43660, GSE182001), and identified 4757 genes (with a FKPM > 1) that were enriched in our dataset (Fig. 8C). We found no distinguishable difference in the gene expression pattern between the two stromal cell activating conditions suggesting LTβ signalling, whether indirectly via B cells or directly via the agonist antibody induced a similar expression profile in eosinophils. Highly enriched IL-33 induced genes (with a FKPM ≥ 5; 3649 transcripts) were analysed using the reactome database to gain insight into the key pathways that were being upregulated in eosinophils (Fig 8D). Reactome pathways that were significantly enriched included the vascular endothelial growth factor (VEGF)-A, VEGFR2 pathway and VEGF signalling (Fig 8D-E), which has been linked to enhanced eosinophil chemotactic and degranulation activity (*37*). Validation by RT-PCR confirmed that *Vegfa* expression was significantly increased in eosinophils that had been co-cultured with LTβR activated FRCs (Fig. 8F). Both gene expression analysis and bulk RNA sequencing showed that eosinophils co-cultured with activated stroma have significantly increased expression of *Icam1* (Fig. 8G) and an enriched expression of adhesion molecules involved in extracellular matrix-receptor interactions and focal adhesion (Fig. S6E-F). This suggested that stromal cell derived IL-33 might enhance eosinophil accumulation within the mLN through enhancing chemotaxis as well as cell-cell adhesion. Additionally, other significantly enriched reactomes included hemostasis, IL-3, IL-5, and GM-CSF signalling and IL-4 and IL-13 signalling (Fig. 8B), which are linked to tissue repair (Fig. 7I), degranulation (Fig. 7J), platelet activation (Fig. S6G) as well as promoting the adaptive immune response through antigen processing and presentation (Fig. S6H), suggesting stroma not only provide a suitable niche for eosinophil recruitment, but support and modulate the eosinophil immune function within the mLN microenvironment.

**Figure 8:**
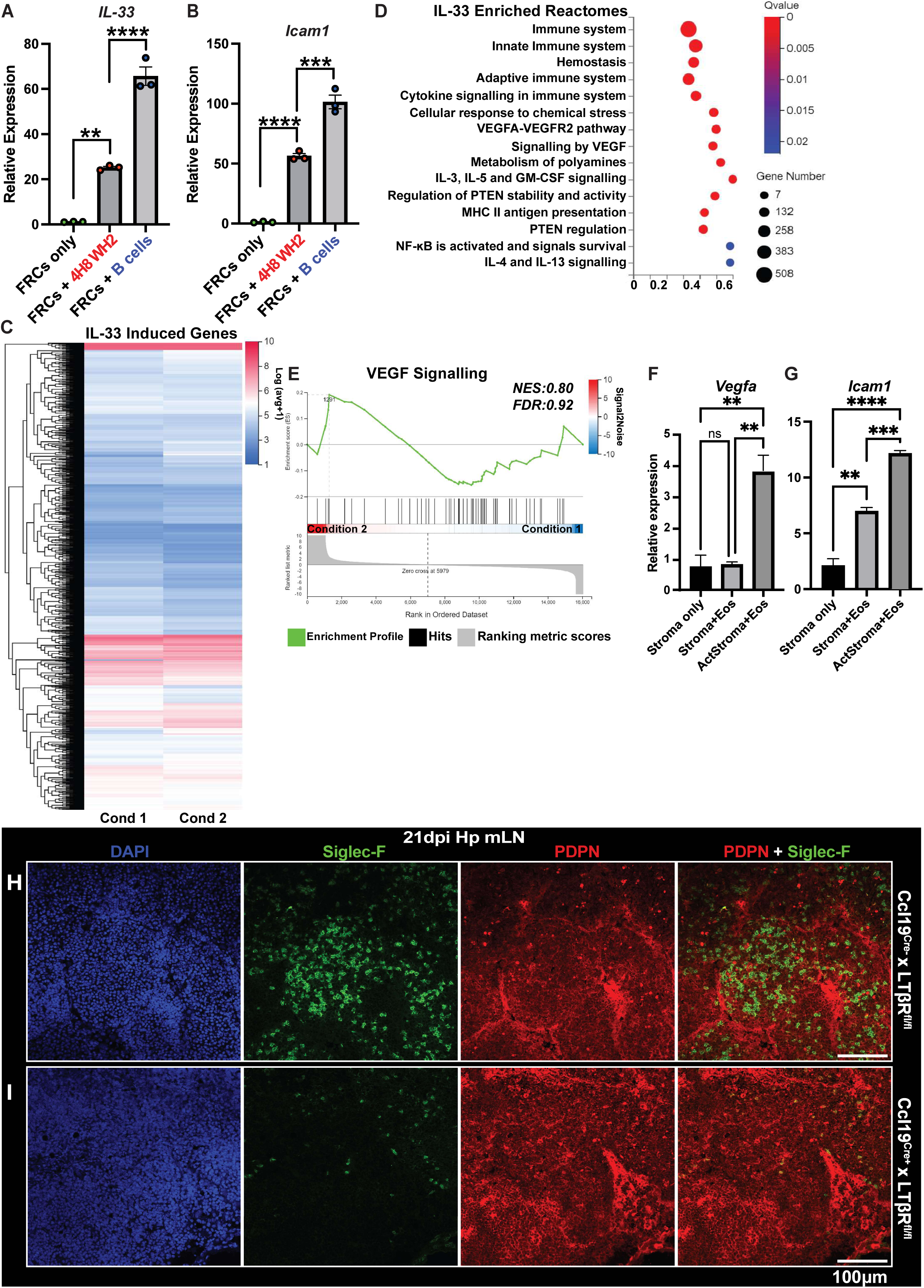
Loss of lymphotoxin-beta receptor (LTβR) on CCL19^cre+^ FRCs attenuates eosinophilia. Activation of FRCs by LTβR ligation or via co-culture with B cells was confirmed by the relative expression of **(A)** *Il33* and **(B)** *Icam1*. Activated stromal cells were co-cultured with eosinophils and analysed using bulk RNA sequencing. **(C)** Heatmap showing IL-33 analysis based on GSE43660 and GSE182001 datasets with an FKPM ≥ 1. **(D)** Reactome bubble chart representing 15 reactome pathways of highly enriched IL-33 induced genes (FKPM ≥ 5). **(E)** GSEA plot and **(F)** *Vegfa* gene expression analysis representing VEGF signalling in eosinophils co-cultured with stromal cells. **(G)** *Icam1* expression in eosinophils co-cultured with LTβR stromal cells. CCL19^cre+ or^ ^Cre-^ x LTβR^fl/fl^ mice were infected with *Hp* and mLN were collected at 21 dpi. **(H-I)** mLN cryosections from 21 dpi mice showing combined immunofluorescence staining for Siglec-F^+^ eosinophils (green), Pdpn^+^ stroma (red) along with DAPI are shown. The data is representative of at least 1 independent experiment and is presented as mean ± SEM. One-way ANOVA tests were used to denote the significance - *p<0.05, **p<0.01 ***p<0.001, ****p<0.0001, ns p>0.05.

As LTβ activation of CCL19^+^ fibroblasts induces cytokine and chemokine secretion (*38*) which we have shown enhances eosinophil activation, survival, and chemotaxis (Fig. 7, 8, Fig. S6), we next aimed to confirm the importance of LTβR ligation of FRCs in supporting eosinophilia into the mLN. Since lymphotoxin expressing B cells are critical for the recruitment of eosinophils (Fig. 6), we hypothesised that the expansion of the lymphoid infrastructure to support incoming eosinophils was governed by LTβR expressing stroma. To assess this, we used mice that have cre-recombinase expression under the control of the CCL19 promoter (*39*) which were crossed to LTβR^fl/fl^ mice resulting in a selective deletion of LTβR on CCL19^+^ stromal cells. *Hp* infection of Cre^+^ (CCL19^Cre+^ x LTβR^fl/fl^) showed a significantly decreased presence of eosinophils within the mLN compared to Cre^-^ (CCL19^Cre-^ x LTβR^fl/fl^) mice (Fig. 8H-I). Thus, our data suggested that FRCs positioned in the mLN interfollicular regions and paracortical regions contribute to eosinophilia into the mLN. Overall, these results highlight the supporting role of FRCs in eosinophil recruitment into the mLN. Our results further highlight the complexity within the lymphoid compartment and open new doors to study potential interactions between eosinophils, immune cells, and stroma in diverse immunological settings (*40–42*).

In summary, by using *Hp* as a model organism we provide a mechanistic view of eosinophil recruitment to the secondary lymphoid organ and highlight a novel role of B cell-stromal cell cross talk in this process (Fig. S7).

## Discussion

The role of stromal cells in eosinophil recruitment to the mLN expands the central paradigm about lymphoid non-hematopoietic stroma which was largely associated with the survival, migration, and function of B cells, T cells, and DCs (*4, 5, 43*). Utilizing Hp as a model organism, we elucidate the intricate mechanism behind the recruitment, sustenance, and functionality of eosinophils within the secondary lymphoid tissues draining the gut.

Our results suggest that helminth-induced mLN stroma activation and remodelling can provide the necessary infrastructure and chemokine expression for the recruitment and retention of eosinophils, which is in line with previous work (*24, 44*) as well as supports the eosinophils-plasma cell association (*30*). The presence of eosinophils in close association with extrafollicular CD138^+^ plasma cells further suggests that they can support the humoral immune response against helminths (*20, 45*). IL-4Rα expression on hematopoietic cells is key to mounting a protective immune response against nematodes (*46, 47*) where IL-4Rα expressing B cells drive the expansion and remodelling of stromal subsets (*9*). This further emphasises the importance of stromal-immune cell interaction towards an enhanced type-2 response by repositioning cells at immune interactive sites (*21, 31*), with previous studies highlighting that immune cell positioning by lymphoid organ stromal cells is key to a rapid T cell response (*48*) as well as B cell survival and function (*4, 21*). As eosinophils have also been implicated in supporting B cell survival (*18*), exploring the cellular constituents at the interfollicular region (IFR) and T/B border, which are important immune niches, will be crucial to determine the spatial cellular diversity in these sites. The IFR lies between the B cell follicles and the T cell zone and is where B cells, T cells, and DCs interact to generate the adaptive immune response. Our results demonstrated that eosinophil recruitment to immune interactive sites within the mLN required IL-4Rα expressing B cells, which supports previous findings (*49*). Our results further confirm the importance of both IL-5 and IL-13 in driving mLN eosinophilia. Both IL-5 and IL-13 play an important role in the type-2 immune response as well as in eosinophil recruitment (*50*) to mucosal and adipose tissues. Our results further suggest that since both FRCs and LECs are capable of producing eosinophilic chemoattractant as well as IL-33 (*51*) that can function as an activator and survival factor for eosinophils (*52, 53*), IL-13 might be dispensable for mLN eosinophilia.

The presence of CCR7 on eosinophils highlights their migratory potential from the periphery to the lymphoid compartment as seen previously in allergic inflammation (*54, 55*). This suggests that stroma-derived eosinophil chemoattractants as well as CCR7 ligands, CCL19 and CCL21, can operate in synergy to localise these granulocytes to distinct regions of the mLN where DC clustering, antigen presentation, and T cell and B cell activation takes place. The localisation of eosinophils in the paracortical region of the mLN is in keeping with the notion that FRCs through the secretion of CCL11, CCL19, and CCL21 (*56*) can recruit eosinophils and supposedly support eosinophil antigen presentation and T-lymphocyte activation (*55*). However, the route and mechanisms of this migration and antigen presentation are not yet well defined and would require further investigation.

The stromal subset expansion mediated through intrinsic B cell IL-4Rα signalling and provision of lymphotoxin further supports the secretion of eosinophil chemoattractant CCL24, which has been previously shown to be an important player in type-2 immunity as well as susceptibility to helminth infections (*57*). These findings support our previous observation and highlight the role of stromal expansion and remodelling towards the development of a type-2 response (*21*). Immgen analysis indicated the differential expression of CCL24 within the stromal subsets, which could be linked to LEC expansion as previously observed (*27*). FRC-B cell crosstalk supports LEC expansion during a helminth infection and loss of LTβR on CCL19^+^ FRCs abrogates LEC expansion and proliferation (*27*); therefore, the reduced eotaxin-2 expression in IL-4Rα^-/-^ mice can be linked, although indirectly, to the lack of LEC expansion. Furthermore, a more detailed analysis of lymphangiogenesis in IL-4Rα^-/-^ mice is required to assess the link between CCL24 expression and LEC expansion. Nevertheless, the reduced LEC numbers seen in infected CCL19Cre^+^ x LTβR^fl/fl^ mice (*27*) directly complement the reduced eosinophilia we observed. A limitation of our study is that we could not compare the contribution of other immune cells such as DCs, macrophages, and ILC2s in driving mLN eosinophilia. Nevertheless, the loss of CCL24 in IL-4Rα^-/-^ mice is in line with previous work highlighting the type-2 dependency of this cytokine in diverse settings (*42, 57, 58*). These results further strengthen the functional role of eosinophils in diverse immunological settings from cancer to thymus regeneration where IL-4Rα plays a key role (*41, 42, 59*).

The remodelling and expansion of MAdCAM1^+^ marginal reticular cells (MRCs) from the subcapsular sinus region to the IFR (*9*) could further support our hypothesis, where α4β7^+^ eosinophil interaction with MAdCAM1^+^ MRCs govern their accumulation at the IFR. Novel genetic tools targeting MAdCAM1^+^ MRCs will need to be developed to reveal the contribution of MRCs in eosinophilia to the mLN.

Eosinophils have been shown to support the survival and function of peripheral B cell populations in both humans and mice (*17*) and stromal cells are known to support B cell survival within the lymphoid organs (*4*). Our findings further strengthen the hypothesis that stromal cells not only regulate local B cell survival but may also indirectly regulate the peripheral population by recruiting eosinophils from the tissue to the mLN which allows these effector cells to migrate back into circulation. The lymphotoxin activated FRC-eosinophil co-culture highlights the capability of activated FRCs to modulate both MHC-II expression as well as ICAM-1 expression in eosinophils. A possible mechanism of such regulation could be that FRC-derived IL-33 directly programs eosinophil ICAM-1 expression (*52*). FRC-derived IL-33 has recently been shown to induce the beiging of white adipose tissue (*60*); a process where eosinophils, ILC2, and type-2 cytokines play a pivotal role (*61*). The upregulation of ICAM-1 on eosinophils and their presence at the IFR of the mLN further strengthen the notion that eosinophils might provide an additional scaffold at these sites for B cell adhesion and synapse formation, as previously reported (*62*). The association of eosinophils with vasculature and FRCs further hints that ICAM-1 supports VEGF-A mediated chemotaxis (*63*), and previous reports have shown that IL-33 induced ICAM-1 expression enhances adhesiveness of eosinophils (*64–66*).

Considering the high levels of IL-4, IL-5, and IL-33 and the constant presence of eosinophils within the mLN over the time course of infection also suggests that the lymphoid microenvironment provides suitable survival niches for granulocytes (*25, 26*). While mLN eosinophilia appears to be dispensable for the anti-helminth response against *Hp* (*47*), our results suggest that the eosinophils contribute to stromal activation and development of the Th2 response. Furthermore, eosinophils could simply pass through the mLN to enter circulation and subsequently migrate to mucosal tissues, like the small intestine, where they regulate tissue homeostasis (*67*) as well as perform effector functions (*68*). However, how eosinophils directly regulate the different stromal subset functions is not fully understood. Although the FRC-eosinophil co-culture provides a suitable *in vitro* system for studying such interaction, future studies with eosinophil-deficient ΔdblGATA mice are needed to deleniate the exact contribution of these cells.

In Summary, our results identify an intricate relationship between B cells and stromal cells in the recruitment of eosinophils to various immune interactive niches within the mLN. IL-4Rα expressing B cells play a key role in this immunological circuit and offer a novel axis to target eosinophilia. Our study further extends the current paradigm about LN stromal cells and highlights that lymphoid stroma can govern the recruitment, migration, and function of not only lymphocytes and myeloid cells but also eosinophils.

## Materials and Methods

### Mice, helminth infection, treatments, and ethical statement

All animal procedures were performed using 7-12-week-old mice (in age and sex-matched groups) in accordance with the institutional Animal Welfare Ethical Review Body (AWERB) and UK Home Office guidelines. All studies were ethically reviewed and approved by an institutional body and carried out under the Animals (Scientific Procedures) Act 1986. C57BL/6J (WT) mice were purchased from Charles River Laboratories. Jht^-/-^, IL-4Ra^-/-^, IL-5^R5/R5^ and IL-13^gfp/gfp^ mice were bred on the C57BL/6J background and maintained under specific pathogen-free conditions. IL-4Rα^-/-^ were maintained at École Polytechnique Fédérale de Lausanne (EPFL), Switzerland (authorization numbers: VD2238.1 and VD3001), and the University of Manchester, United Kingdom. IL-5^R5/R5^ and IL-13^gfp/gfp^ mice were housed and maintained at the Univeristy of Manchester. LT-β^−/−^ and TCRβδ^−/−^ mice were maintained at the University of Lausanne, Switzerland, and were a kind gift from Sanjiv A. Luther. CCL19^Cre^x LTβR^fl/fl^ mice were provided by Burkhard Ludewig. All mice were maintained under specific pathogen-free conditions, with water and chow provided ad libitum. Throughout the study, mice were orally infected with 200 L3 stage *Heligmosomoides polygyrus (Hp)* and were sacrificed at indicated time points to study the mesenteric lymph node (mLN).

### Bone Marrow Chimera

Bone marrow chimeras were set up as previously described (*27*). In brief, bone marrow (BM) was obtained from the femur and tibia of donor mice and injected intravenously into C57BL/6J or IL-4Rα^-/-^ recipient mice that had been previously irradiated twice with 450 rad with 4-hour intervals between irradiation sessions. All mice were maintained in specific pathogen-free conditions. For the generation of complete BM chimeric mice which lacked IL-4Rα on CD45^-^ or CD45^+^ cells, IL-4Rα^-/-^ mice received WT-BM or WT mice received IL-4Rα^-/-^ BM, respectively. Mice lacking IL-4Rα on B cells and WT chimeras were generated using irradiated WT mice reconstituted with 80% Jht^-/-^ BM and 20% IL-4Rα^-/-^ BM or with 80% Jht^-/-^ and 20% WT BM, respectively. For the generation of mice with B cells or T cells lacking lymphotoxin expression (B-Ltβ^−/−^ or T-Ltβ^−/−^), C57BL/6J recipient mice were reconstituted with 80% Jht^−/−^ or 80% TCRβδ^−/−^ bone marrow plus 20% Ltβ^−/−^ bone marrow. Control mice were generated using WT recipient mice which were reconstituted with 80% Jht^−/−^ or 80% TCRβδ^−/−^ bone marrow plus 20% WT bone marrow, respectively. All chimeric mice received an antibiotic in their drinking water for 4-6 weeks following the BM reconstitution and were subjected to infection 8 weeks following reconstitution.

### Flow cytometry

At the given experimental time points, mice were killed and the whole mLNs were isolated, and single-cell suspensions were generated using enzymatic digestion. mLNs were incubated at 37°C in digestion medium (RPMI-1640 supplemented with 0.8 mg/mL dispase, 0.2 mg/mL Collagenase P [Roche] and 0.1 mg/mL DNase I [Invitrogen]) and gently pipetted at 5-minute intervals to ensure complete dissociation of the tissue. Following digestion, the single-cell suspension was filtered through a 70 μm cell strainer, counted, and resuspended in FACS buffer (PBS supplemented with 2% FBS and 5 mM EDTA). For eosinophil staining, cells were incubated for 30 minutes with an antibody cocktail and identified as CD45^+^CD3^-^CD19^-^ CD11c^-^Ly6G^-^CD11b^+^Siglec-F^+^ cells (Fig. S1). Samples were acquired on the BD machine and analysed using FlowJo V.10.

### Histology and immunofluorescence microscopy

The entire mLN chain was carefully dissected, weighed, imaged, and embedded in Tissue-Tek optimum cutting temperature control compound (OCT) and frozen in an ethanol ice bath. Cryostat sections (8 mm in thickness) were cut from from a 400 μm span of the mLN onto Superfrost Plus glass slides, air-dried, and fixed for 10-15 minutes in ice-cold acetone. After rehydration in PBS, sections were blocked with 1% (w/v) BSA and 1-4% (v/v) mouse and donkey serum. Immunofluorescence staining was performed using antibodies (Siglec-F, Pdpn, Lyve1, CD138, etc.). Sections were incubated overnight at 4°C in the primary antibodies. On the following day, sections were washed four times with PBS, and primary antibodies were detected using fluorescently labelled secondary antibodies and nuclei counterstained with DAPI prior to mounting sections using ProLong anti-fade reagents (Life Technologies). Images were acquired on an Olympus VS120−SL full slide scanner using a 20x/0.75 air objective and an OlympusXM10 B/W camera or with Nanozoomer Slide-scanner, Hamamatsu equipped with fluorescence filters for DAPI, SpectrumAqua, FITC, TRITC, Texas Red and Cy5 with 20x objective. On occasions, LSM710 (20x/1.2 or 40x/1.4 using regular PMTs, 1AU pinhole, Z spacing 1−10um) laser scanning confocal microscope was used to acquire images for quantification. For quantitative measurements, immunofluorescence images from naïve and infected mice mLN were acquired and segmented using an ImageJ/Fiji pipeline. A threshold−based approach was used to measure areas specific for a given marker and expressed as the percentage of total area within the tissue occupied by the given marker. The final figure panels (graphs and images) were arranged and converted to .TIF file (LZW compression) using Adobe Photoshop 24.02 edition 2023.

### Deep tissue imaging of mLN to visualise eosinophils

C57BL/6J and IL-4Rα**^−/−^**mice were infected with *Hp* and the mLN was collected at 21 dpi for vibratome sectioning and deep tissue imaging to identify eosinophil association with stromal cells. 200μm thick vibratome sections were generated using Leica VT1200S vibratome. Post sectioning, tissue was blocked in blocking solution followed by two stage staining as described previously with slight modification (*27*). We used antibodies against well-established markers like Siglec-F, Lyve1, and GP38 to identify eosinophils, lymphatics, and FRCs, respectively. The stained tissue was cleared using the Miltenyi Biotec MACS Clearing Kit protocol as per the manufacturer’s recommendation. Stained samples were imaged using a Zeiss confocal microscope. The 3D reconstruction and movies were made using IMARIS (Bitplane) and exported as a .mpg file.

### Eosinophil-stromal cell co-culture

Bone marrow-derived eosinophils were generated *in vitro* as described previously (*42*). Briefly, bone marrow cells were obtained from femurs and tibiae of WT C57BL/6 mice by flushing with complete RPMI 1640 media containing 10% heat inactivated fetal bovine serum (FBS) (Sigma-Aldrich, non-USA origin). Red blood cells were lysed using ACK lysis buffer, and 1 × 10^6^ cells/mL were plated in RPMI-1640 medium containing 10% FBS, penicillin-streptomycin and further supplemented with stem cell factor (100 ng/ml), and FLT3Ligand (100 ng/ml) (both from Miltenyi Biotec) from culture days 0 to 4. Cultures were changed on day 4 to new complete RPMI 1640 medium containing recombinant murine IL-5 (10 ng/ml, Peprotech) and further fed with IL-5 every other day between days 10 to 14. Cells were collected on day 14 (typically containing more than 92% eosinophils), counted, and used for the co-culture experiment. Naïve B-cells were isolated from whole mLN by negative selection as described previously (*27*). mLN derived stromal cells were cultured i*n vitro* as previously described (*69*). In brief, single-cell suspensions from whole mLN were generated using enzymatic digestion and plated at a density of 20 x 10^6^ in a 6-well plate. After 24 hours non-adherent cells were removed and adherent cells were further cultured for 7-10 days, with culture media changed every other day. After 7-10 days of culture, the adherent cells containing primarily FRCs and LECs were removed and seeded in a 24-well plate at a density of 0.5 x 10^6^ per well. Stromal cells were co-cultured with 1 x 10^6^ naïve B cells or stimulated with 2 µg/mL anti-LTβR antibody for 16 hours in triplicate wells. The stromal cells were washed thrice with PBS to remove agonist antibody or B cells before adding the eosinophils. After washing 0.5 x 10^6^ eosinophils were added to activated stroma and co-cultured for an additional 8 hours. Post co-culture, eosinophils from triplicate wells were pooled and stroma were harvested and stored in RNA lysis buffer at -80°C until analysed for gene expression and bulk RNA sequencing as detailed below. In a separate set of experiments, FRCs were either stimulated with anti-LTβR agonist antibody (Clone 4H8WH2, 2μg/ml, Adipogen) or with B cells isolated from mLN using negative selection (1:5 ratio) for 24 hours. Post stimulation, cells were washed, and triplicate wells were pooled and stored in RNA lysis buffer at -80°C until analysed for gene expression.

### RNA isolation and qRT-PCR analysis

The stromal and cellular fractions were separated as previously described (*70*). In brief, the mLN was gently mashed through a 40μm cell strainer using a 5 mL syringe plunger. The filtered cells represented the cellular fraction, and the matter left on the strainer represented the stromal cell fraction. The RNA from the stromal fraction was extracted with Direct-zol RNA MiniPrep Kit (Zymo Research) and reverse transcribed using RevertAid cDNA synthesis reagents (Thermo Scientific) for qPCR analysis. RNA from the co-culture experiment was extracted using the RNeasy UCP Micro Kit (Qiagen), and reverse transcribed using a high-capacity cDNA reverse transcription kit (Applied Biosystems). The qPCR was performed using PowerUp SYBR Green Master Mix (Applied Biosystems) on an Applied Biosystems 7900HT system. The following primers were used to detect the various gene expression. *Ccl5-Fw*: CCTCACCATCATCCTCACTGCA, *Ccl5-Rv*: TCTTCTCTGGGTTGGCACACAC; *Ccl11-Fw*: CCCAACACACTACTGAAGAGCTACAA *Ccl11-Rv*: TTTGCCCAACCTGGTCTTG; *Ccl24-Fw*: GCAGCATCTGTCCCAAGG, *Ccl24-Rv*: GCAGCTTGGGGTCAGTACA; *Il33-Fw*: CACATTGAGCATCCAAGGAA; *Il33-Rv*: ACAGATTGGTCATTGTATGTACTCAG; *Vegfa-Fw*: GCTGTACCTCCACCATGCCAAG; *Vegfa-Rv*: ACTCCAGGGCTTCATCG; *Icam1-Fw*: GACAGTACTGTACCACTCTC; *Icam1*-*Rv*: CCTGAGCCTTCTGTAACTTG; *MhcII-Fw*: CTCCGAAAGGCATTTCGT; *MhcII-Rv*: CTGGCTGTTCCAGTACTC; *Ccr7-Fw:* AGAGGCTCAAGACCATGACGGA ; *Ccr7-Rv*: TCCAGGACTTGGCTTCGCTGTA; *Il6-Fw*: GCTACCAAACTGGATATAATCAGGA; *Il6-Rv*: CCAGGTAGCTATGG-TACTCCAGAA; *Il1*β*-Fw*: CAGTTGTCTAATGGGAACGTCA; *Il1*β*-Rv*: GCACCTTCTTTTCCTTCATCTTT; Ccl20-Fw: GTGGGTTTCACAAGACAGATGGC; Ccl20-Rv: CCAGTTCTGCTTTGGATCAGCG; *Gapdh-Fw*: GTGCCAGCCTCGTCCCG, *Gapdh-Rv*: TTGCCGTGAGTGGAGTCA; *β-actin-Fw*: CTTTTCACGGTTGGCCTTAG, *β-actin-Rv*: CCCTGAAGTACCCCATTGAAC. Gene expression was normalised against endogenous control and 2^-^δδct values were calculated and presented as relative expression to naïve cells. The eosinophil chemoattractant profile for lymphoid stromal cells were also analysed using datasets available on Immgen.org. The data were presented as expression values normalised by DESeq2 without any modification.

### Bulk RNA sequencing and analysis

Experiments were performed in triplicate wells and eosinophils were pooled post stimulation to create a single sample. RNAseq was performed for duplicate samples per condition. Total RNA from eosinophils isolated from the co-culture was extracted using the RNeasy UCP Micro Kit (Qiagen) according to the manufacturer’s instructions. cDNA libraries were constructed and sequenced by BGI Genomics (China) using DNBSEQ sequencing technology. Raw transcripts were filtered through SOAPnuke (V1.5.2) and hierarchical indexing for spliced alignment of transcripts 2 (HISAT2, V2.0.4) software was used to map the raw data reads. Raw data reads were mapped to the murine reference genome *Mus musculus*, NCIB: GCF_000001635.27_GRCm39. NOISeq analysis was used to identify differentially expressed genes (DEGs). DEGs were defined as log2 of the sample (expression value + 1). Data were directly deposited to the BGI-Dr Tom web-based analysis portal. Read counts were normalised to FKPM (fragments per kilobase of transcript per million mapped reads). Gene set enrichment analysis (GSEA) was performed using normalised read counts, with a max threshold of 500 and a minimum threshold of 15 reads. GSEA enrichment plots were generated using the KeggPathway database. Phyper function in R was used to determine the P-value and the Q-value was obtained by correcting the false discovery rate (FRD) of the p-value. All data was analysed and Reactome pathway enrichment was carried out using the BGI Dr. Tom system. All heatmaps, GSEA graphs, and enrichment plots were exported as .png files.

### STRING analysis

Enriched genes from the bulk RNA seq were mined and analysed using STRING platform V12 (https://cn.string-db.org/) and Cytoscape V3.10.1 (https://cytoscape.org/index.html). The species was restricted to *Mus musculus* and gene IDs derived from Dr. Tom were uploaded to the platforms. Using the STRING platform subnetwork analysis was performed on a given set of genes using GO term and biological processes annotation. The known interactions were shown using cyan and magenta lines and the predicted interactions were represented using green, red, blue, light green, and black lines. Kmeans clustering was also performed to find two defined clusters of proteins within the regulation of IL-5 production GO network, with each cluster highlighted using red or green colour. The Cytoscape platform was used to highlight the gene expression (FKPM) of each individual gene within a given network. The strength of the interaction is depicted by the line density based on predicted protein interactions from the String: protein query plug-in. The network maps were directly exported as a .png file. The geneset analysis datasets were exported to excel files and both counts within the network as well as the total network and background number were presented.

### Statistical Analysis

Flow cytometry and gene expression analyses are expressed as means ± SEM. Statistical analyses were performed using a non-parametric Mann-Whitney Student’s T-test and ANOVA as indicated with post-hoc tests (Bonferronis multicomparison test). p-values indicated as p<0.05 (*), p<0.01 (**), p<0.001 (***), p<0.0001 (****). Graph generation and statistical analyses were performed using Prism version 9 software (Graph pad, La Jolla, CA).

## Data availability

All relevant data supporting the research findings of this study are available within the paper or provided within the supplementary files.

## Author contributions

Conceptualisation and writing (original draft), E.B. and L.K.D. Methodology, investigation, analysis, and validation, E.B., R.F., N.L.H., M.R.H., L.K.D; Resources, L.K.D, M.R.H., N.L.H., L.K.J., B.L; editing of the draft, all authors; overall project conception, supervision, and funding, L.K.D.

## Acknowledgements

L.K.D is supported by the Barts Charity-Rising Star fellow program (Grant code MICG1E1R, to L.K.D). M.R.H. is supported by a Sir Henry Dale Fellowship jointly funded by the Wellcome Trust and the Royal Society (Grant Number 105644/Z/14/Z). R.F. is funded by a Wellcome Trust by the Immunomatrix in Complex Disease PhD programme at the University of Manchester. A special thanks to Olivier Burri and Arne Seitz from the Bio-Imaging and Optics Platform, EPFL, Switzerland for ImageJ/Fiji tools and advice for slide scanner image analysis. We thank Biogen Idec for providing reagents for lymphotoxin studies and Elke Scandella, Kantonsspital, and St. Gallen for genotyping CCL19^Cre^x LTβR^fl/fl^ mice. We also thank Sanjiv A. Luther, Department of Immunobiology, University of Lausanne, for providing LT-β^−/−^ and TCRβδ^−/−^ mice bone marrows and Premkumar Palanisamy, William Harvey Research Institute, Queen Mary University of London, for assistance with animal husbandry.

## Competing financial interests

The authors declare no competing financial interests.

## Materials & Correspondence

All correspondence and material requests should be addressed to Dr. Lalit Kumar Dubey.

---

**Supplementary Figure 1: Characterisation of mLN eosinophils post helminth infection**

C57BL/6J mice were infected with *Hp* and the entire chain of the mLN was collected at 0 (naïve), 7, 14, and 21 dpi and analysed using flow cytometry. (A) The pseudocolour dot plots show the gating strategy for the mLN eosinophils. Eosinophils were identified as CD45^+^CD3^-^ CD19^-^CD11c^-^Ly6G^-^CD11b^+^Siglec-F^+^ cells. (B) The time course analysis of mLN showing eosinophil accumulation within the mLN. (C) mLN total weight and (D) total cell count in naïve and 21 dpi WT and IL-4Rα^-/-^ mice. IL-13^gfp/gfp^ and IL-5^Red/R5^ mice were infected with *Hp* and the entire chain of the mLN was collected at 0 (naïve) and 21 dpi and analysed by flow cytometry. (E) Percentage and (F) absolute number of CD45^+^CD11b^+^SiglecF^+^ cells in IL-13^gfp/gfp^ mice, and (G) percentage and (H) absolute number of CD45^+^CD11b^+^SiglecF^+^ cells in IL-5^Red/R5^ mice. Data represent mean ± SEM and are representative of two independent experiments with n ≥3-4 mice per group.

**Supplementary Figure 2: Visualisation of mLN eosinophils post helminth infection**

C57BL/6J (WT) and IL-4Rα knockout (IL-4Ra^−/−^) mice were either left naïve or infected with *Hp* and the mLN was collected at 21 dpi. (A-B) mLN cryosections showing combined staining for the eosinophils (Siglec-F^+^, green) and lymphatic endothelial cells (LECs, Lyve1^+,^ red) to highlight the medullary region in WT and IL-4Rα^−/−^ mice during homeostasis. (C) mLN cryosections showing combined staining for the DAPI (blue), eosinophils (Siglec-F^+^, green), and LECs (Lyve1^+^, red) cells in WT and IL-4Rα^−/−^ mice post *Hp* infection. The white insets represent eosinophil accumulation at the B/T border. Scale bar = 200μm or 100 μm. Images are representative of ≥5 different experiments with n ≥2-3 mice/group/time point.

**Supplementary Figure 3: Plasma cells co-localise with lymphatics in an IL-4Rα dependent manner**

C57BL/6J and IL-4Rα ^−/−^ mice were infected with *Hp* and the entire chain of the mLN collected at 0 (naïve) and 21 dpi. (A) Naïve and (B) infected WT and IL-4Rα^−/−^ mLN cryosections showing combined staining for the CD138^+^ plasma cells (purple) and LECs (Lyve1^+^, brown). Scale bar = 200μm. Images are representative of ≥3 different experiments with n ≥2-3 mice/group/time point

**Supplementary Figure 4: Visualisation of mLN eosinophilia during homeostasis in mice lacking IL-4Rα on B cells**

Complete bone marrow chimera mice (donor strain/recipient strain) showing mLN eosinophilia during helminth infection. (A) IL-4Rα^−/−^ mice received C57BL/6J (WT) bone marrow and (B) C57BL/6J mice received IL-4Rα^−/−^ mice bone marrow. The resulting animals lacked IL-4Rα on (A)CD45^-^ and (B) CD45^+^ cells. (C-D) mLN cryosections from chimeric naïve mice showing immunofluorescence image staining for Lyve1^+^ (white), Pdpn (red), and Siglec-F^+^ (green) are shown. Mixed bone marrow chimeras were generated using lethally irradiated wildtype (WT) recipients reconstituted with (E) bone marrow cells from B cell deficient (Jht^−/−^) mice mixed with bone marrow cells from IL-4Rα^−/−^ mice (Jht^−/−^ + IL-4Rα^−/−^). (F) Control mice received B cell deficient bone marrow mixed with WT cells (Jht^−/−^ + WT). (G-H) mLN cryosections from Jht^−/−^ + IL-4Rα^−/−^ and control mixed bone marrow chimeric naïve mice showing immunofluorescence image staining for Lyve1^+^ (white), Pdpn^+^ (red) and Siglec-F^+^ (green). Scale bar = 200 µm. The images are from representative mice and from two independent experiments with n ≥2-4 mice per group.

**Supplementary Figure 5: Visualisation of mLN eosinophils during homeostasis in mice lacking lymphotoxin expressing B cells or T cells**

Mixed bone marrow chimeras were generated as described in the methods section. Chimeric mice lacking lymphotoxin expression distinctly on either (A) B cells (Jht^−/−^ + LTβ^−/−^) or (C) T cells (TCRδβ^−/−^ + LTβ^−/−^) were compared to respective control mice having (B) B-cell or (D) T-cell sufficient for lymphotoxin. All chimeric mice mLN were collected at 21 dpi. Chimeric mice lacking lymphotoxin on (A) B cells (Jht^−/−^ + LTβ^−/−^) and their respective (B) controls (Jht^−/−^ + WT) immunofluorescence staining for Pdpn (blue), Siglec-F^+^ (yellow) and Lyve1^+^ (magenta). Chimeric mice lacking lymphotoxin on (C) T cells (TCRδβ^−/−^ + LTβ^−/−^) and respective (D) controls (TCRδβ^−/−^ + WT) showing eosinophilia (yellow) and LECs (magenta). Scale bar = 200 μm.

**Supplementary Figure 6: Co-culturing eosinophils with activated FRCs induces an activated phenotype in eosinophils**

Activated FRCs were co-cultured with eosinophils and analysed by bulk RNA sequencing. Heatmaps showing the expression of (A) cytokines, (B) chemokines, and (C) immunoglobulin receptors. (D) STRING analysis showing the complete network in the regulation of IL-5 production gene ontology (GO) pathway (GO:0032674), showing 2 Kmeans clusters. Known interactions are indicated by cyan (curated database) and magenta (experimentally determined), and predicted interactions are indicated by black (co-expression), green (textmining), and dark blue (gene co-occurrence). GSEA plots showing enrichment of genes in (E) ECM-receptor interactions, (F) focal adhesion (G) platelet activation, and (H) antigen processing and presentation.

**Supplementary Figure 7: Graphical summary**

Graphical summary highlighting the mechanism of eosinophil accumulation within the mLN. An enriched type-2 environment where IL-4Rα driven activation of B cells promotes an enhanced B cell-stromal interaction leading to eotaxin expression which drives eosinophil recruitment into the mLN cortical/paracortical and interfollicular regions. Loss of IL-4Rα or lymphotoxin expression on B cells attenuates the cross talk with LTβR expressing lymphoid stromal cells thus leading to reduced chemoattractant availability leading to reduced eosinophilia within the mLN.

**Supplementary movie 1:**

3D view of a vibratome section showing combined immunofluorescence staining for eosinophils (Siglec-F+; green), LECs (Lyve1; red), and Pdpn^+^ stroma (blue) from mLN of 21 dpi C57BL/6J (WT) mice.

**Supplementary movie 2:**

3D view of a vibratome section showing combined immunofluorescence staining for eosinophils (Siglec-F+; green), LECs (Lyve1; red), and Pdpn^+^ stroma (blue) from mLN of 21 dpi IL-4Ra^−/−^ mice.

## Notes

### Competing Interest Statement

The authors have declared no competing interest.

