## Supplementary figures and images for "B cell-stromal cell cross talk drives mesenteric lymph node eosinophilia during intestinal helminth infection"

### Supplemental Figure-1

#### Supplementary Figure-1

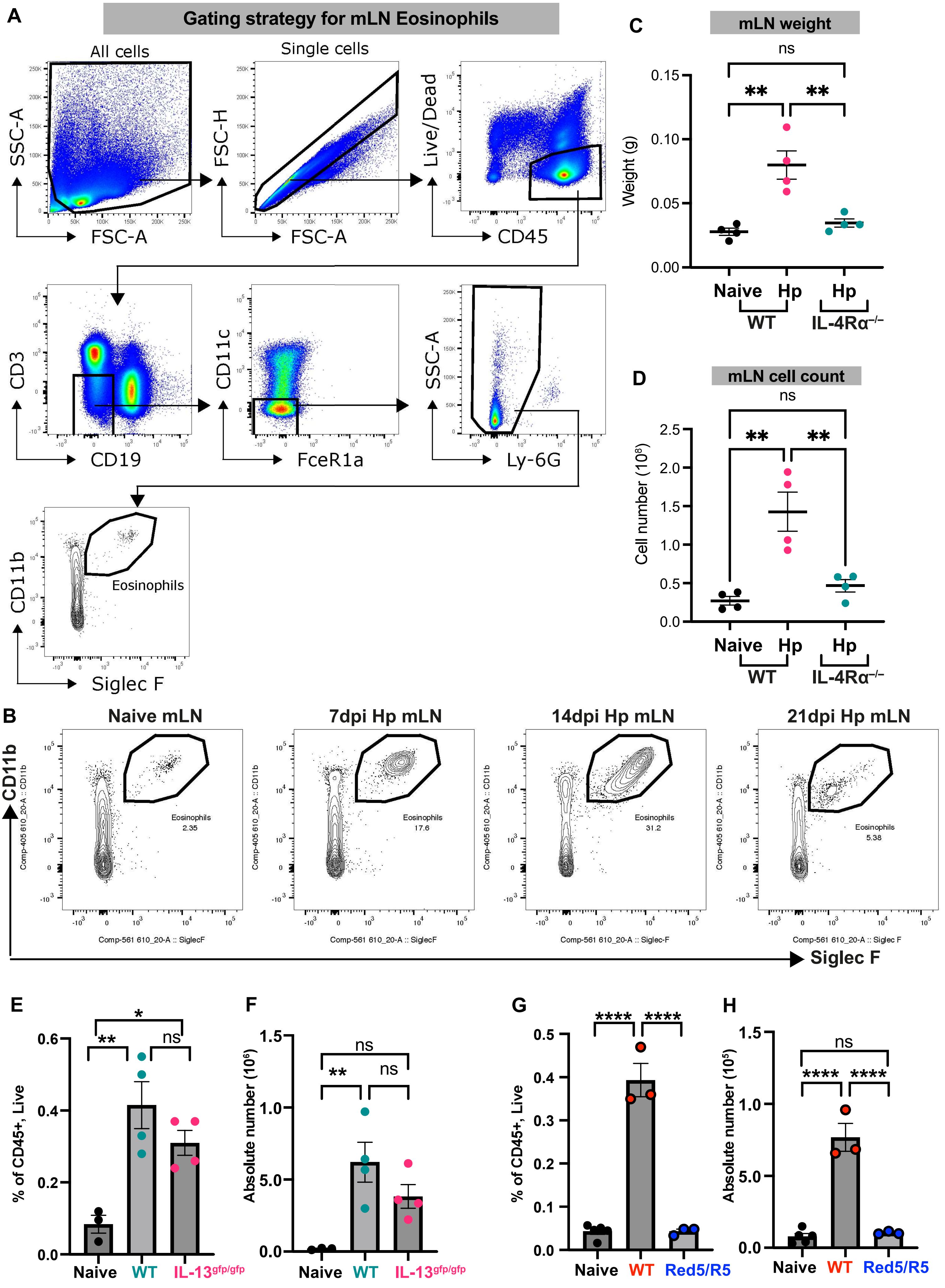

### Supplemental Figure-2

#### Supplementary Figure-2

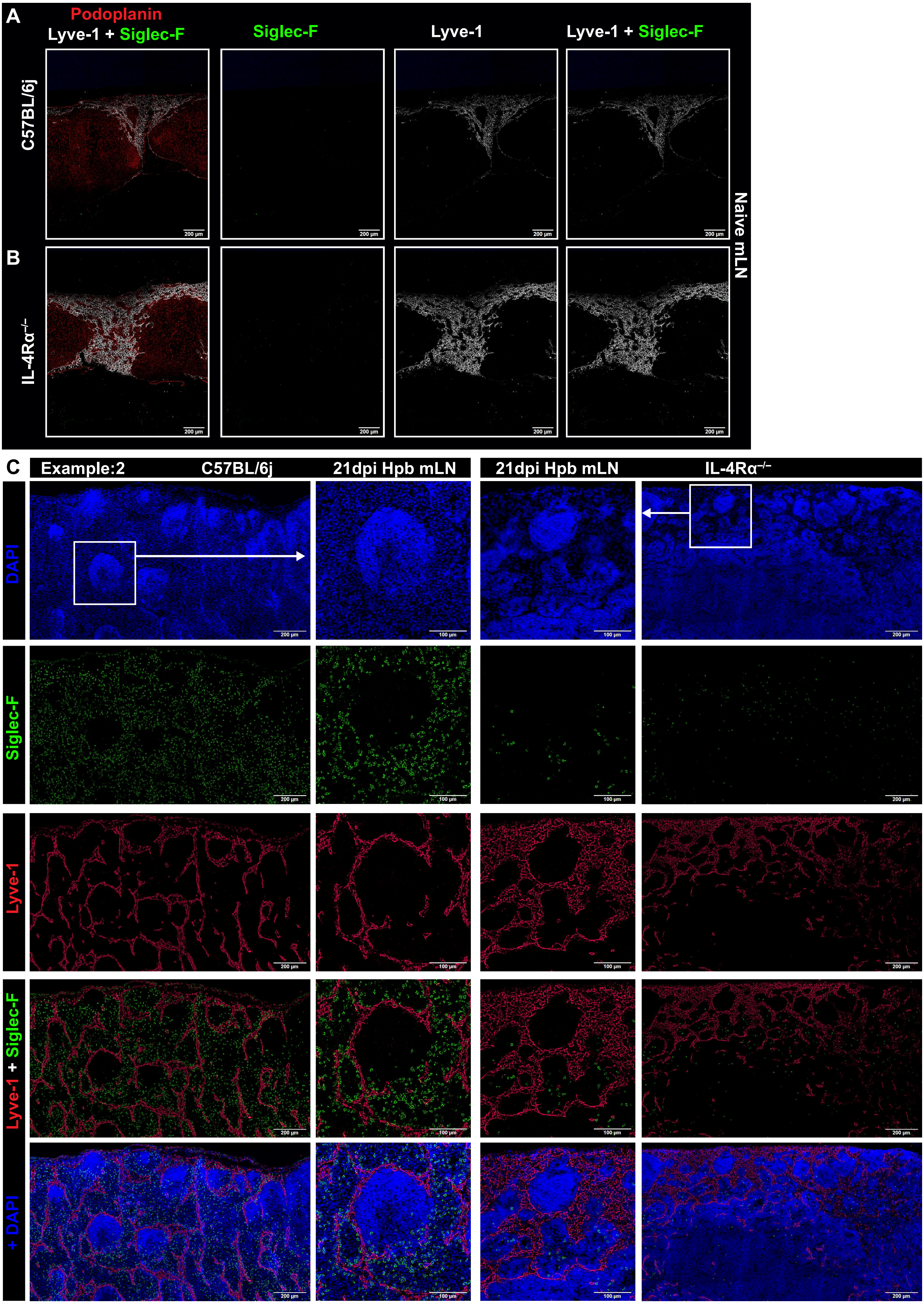

### Supplemental Figure-3

Supplementary Figure-3

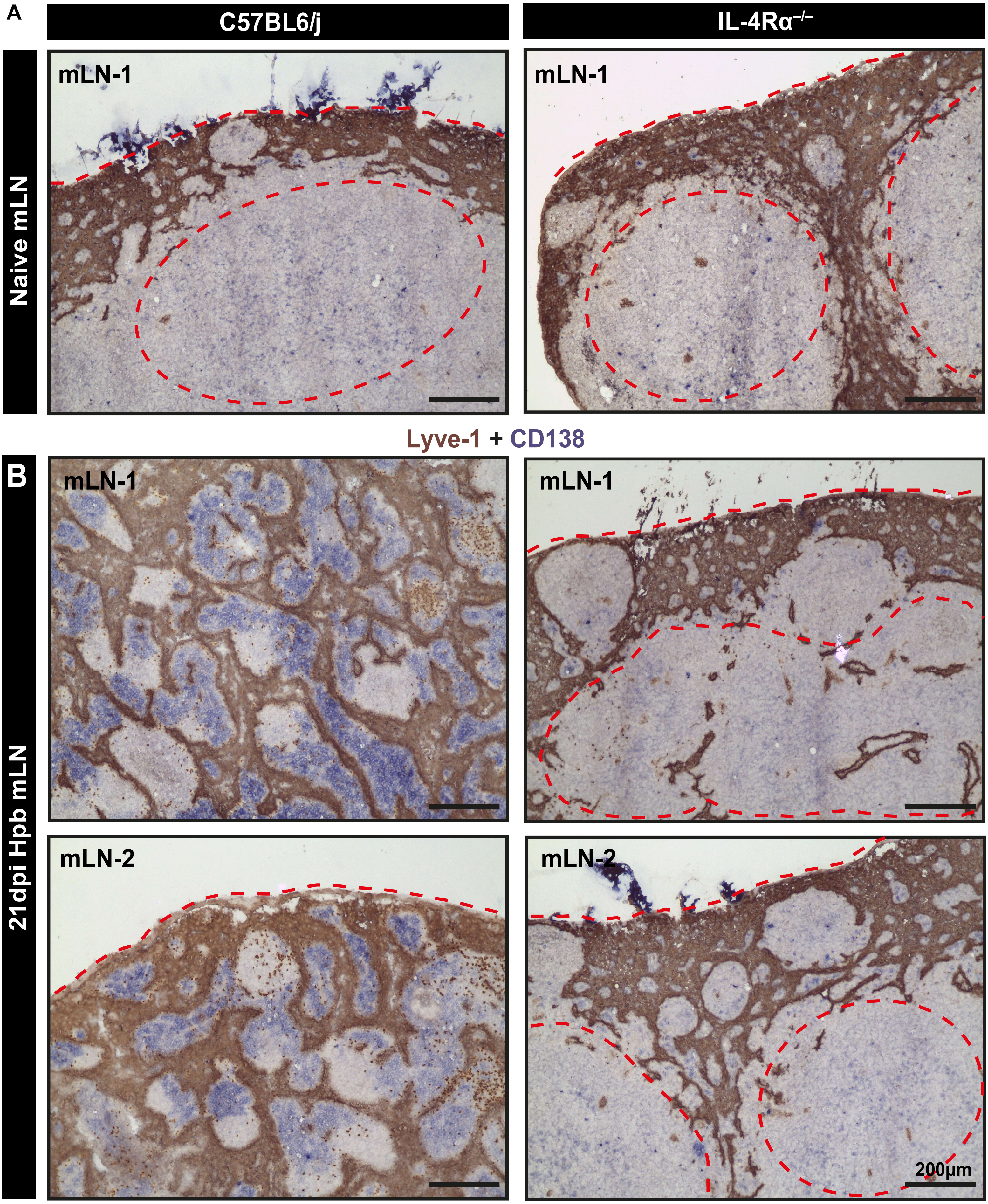

### Supplemental Figure-4

#### Supplementary Figure-4

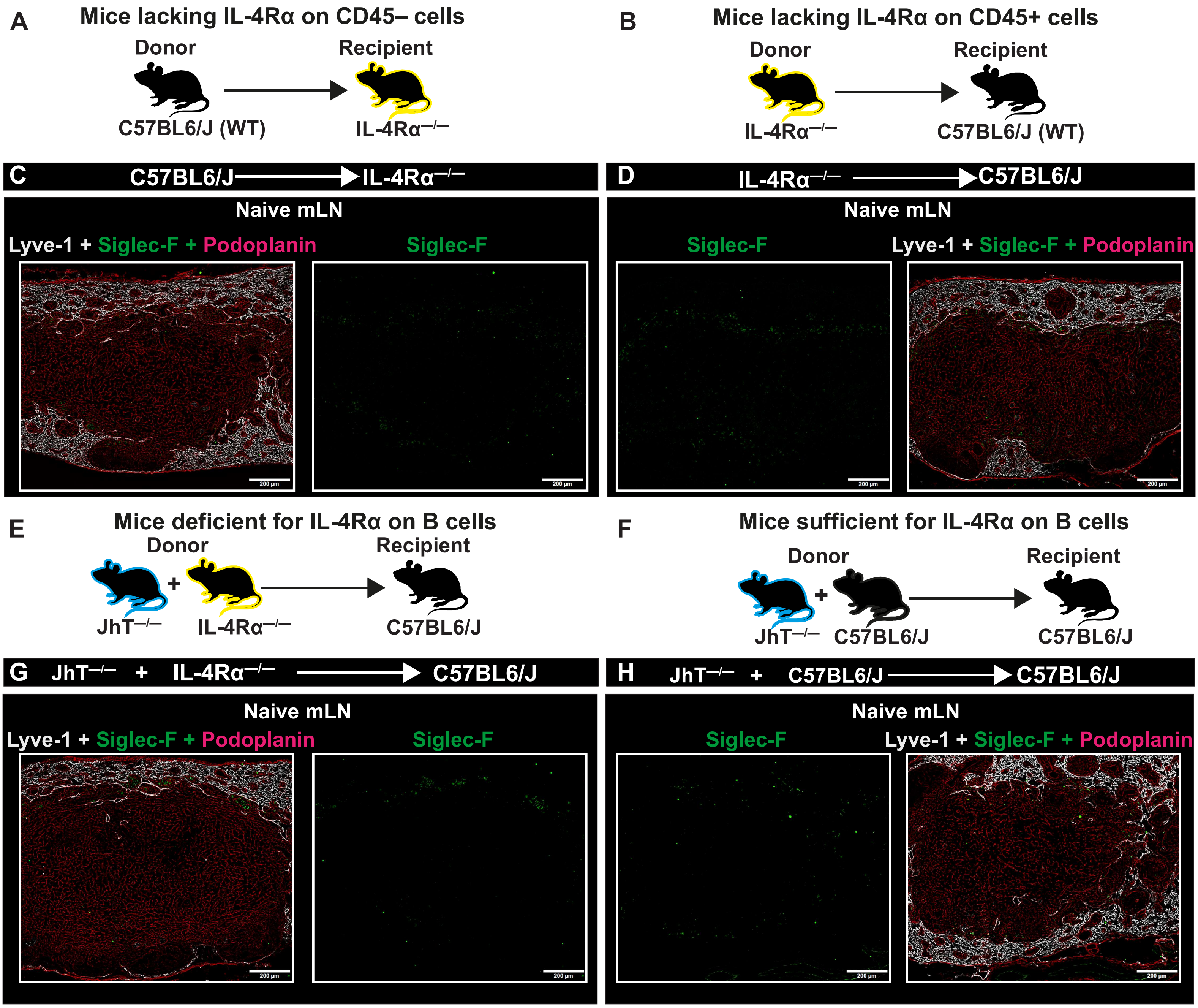

### Supplemental Figure-5

Supplementary Figure-5

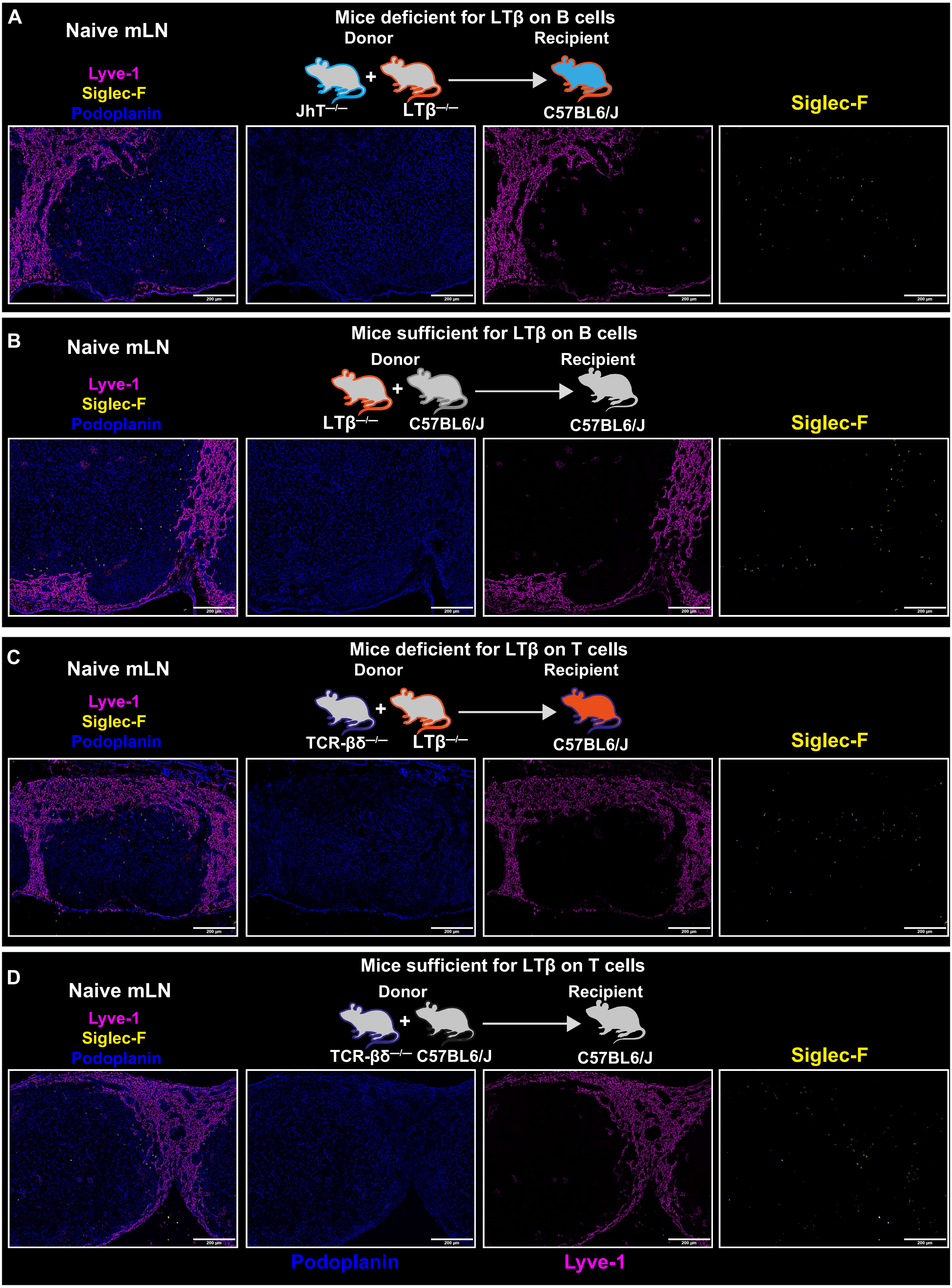

### Supplemental Figure-6

Supplementary Figure-6

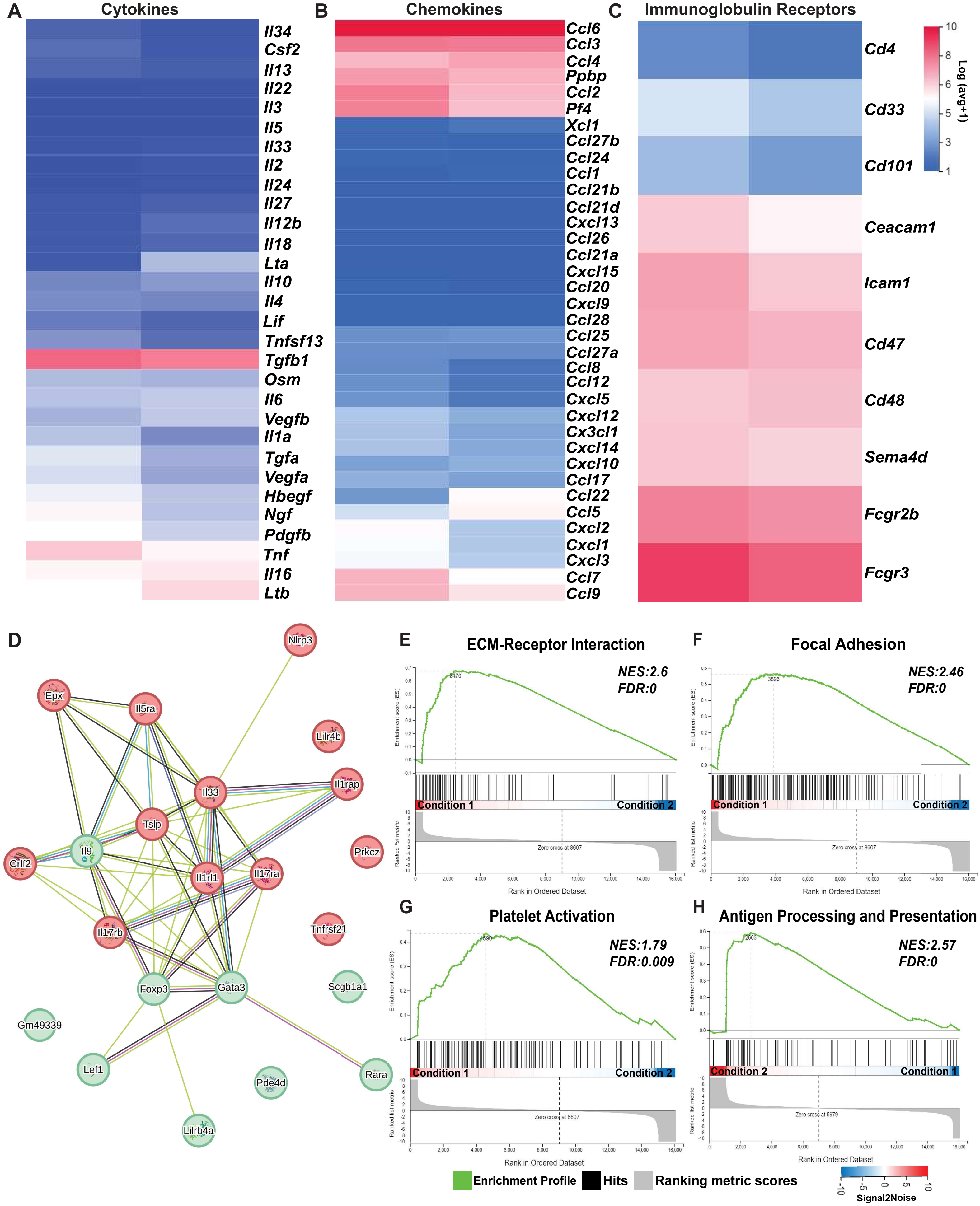

### Supplemental Figure-7

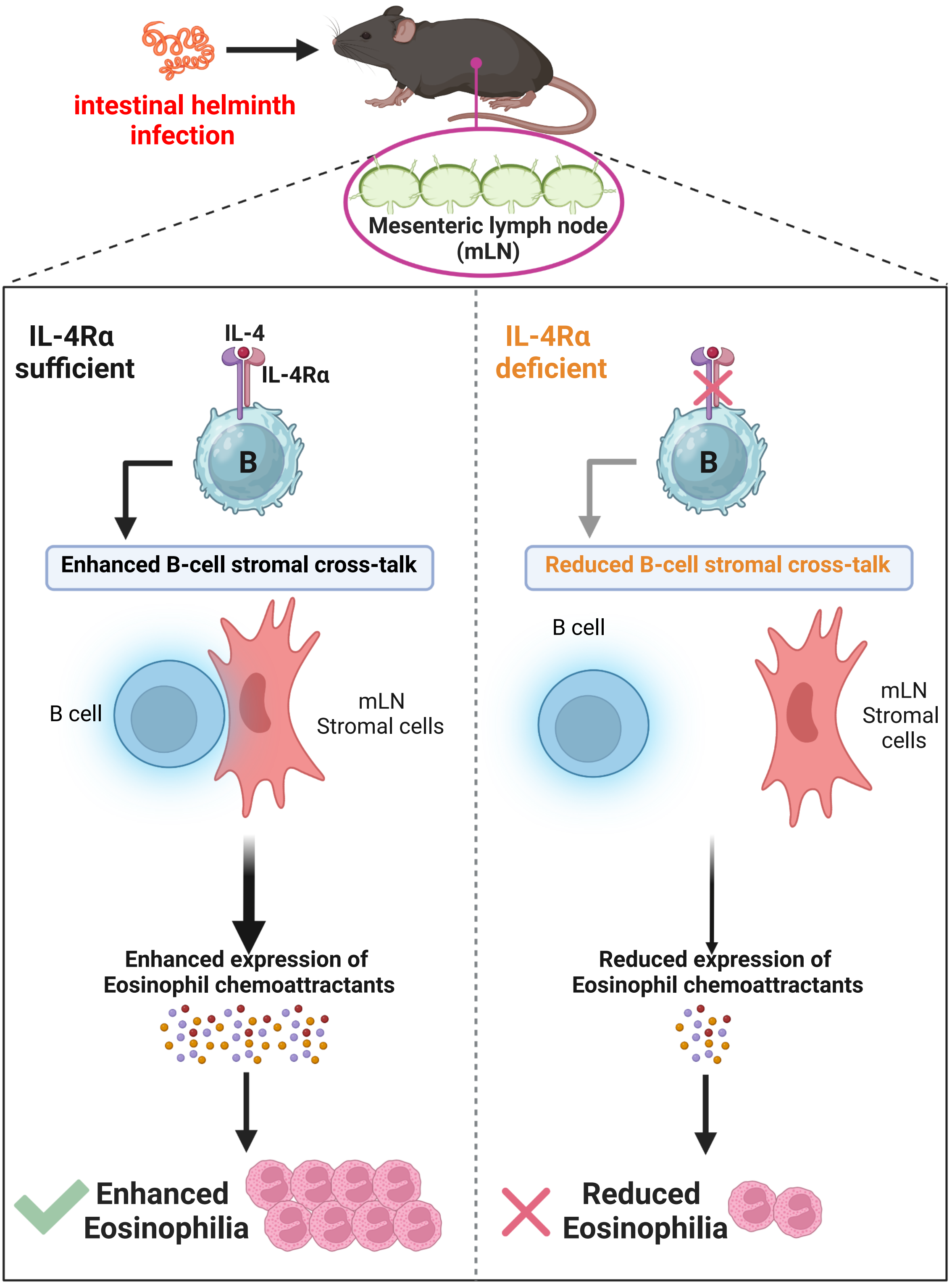
